# Histone Fold mediated heterodimerization specifies the selective association of TAF12 paralogs with TFIID and SAGA complexes in *Candida albicans*

**DOI:** 10.64898/2026.02.06.704445

**Authors:** Poonam Poonia, Krishnamurthy Natarajan

## Abstract

Transcription initiation in eukaryotes is coordinated by the multisubunit coactivator complexes TFIID and SAGA, which share five core TBP-associated factors (TAFs) that assemble through histone fold (HF) mediated heterodimerization. While, in most organisms, a single TAF12 incorporates in both complexes; however, *Candida albicans* uniquely encodes two *TAF12* paralogs, *TAF12* and *TAF12L*, which associate preferentially with TFIID and SAGA, respectively. The molecular basis and functional consequences of this specialization remain unclear. Here, we demonstrate that Taf12 and Taf12L are functionally non-redundant and show strict complex specificity in vivo, even under conditions where the alternate paralog is depleted. Taf12 associates exclusively with TFIID through Taf4, whereas Taf12L incorporates specifically into SAGA through Ada1. Ectopic expression experiments reveal limited and asymmetric cross-complementation, wherein Taf12L led to partial growth rescue and incorporation into TFIID in absence of Taf12 but not vice versa. Biochemical and genetic analyses further show that the conserved histone fold domains (HFDs) of both paralogs are sufficient for biological function and complex incorporation. In vitro interaction assays uncover intrinsic differences in binding selectivity of Histone fold domains, with HFD-Taf12L displaying strong preference for Ada1, while HFD-Taf12 exhibits more promiscuous binding. Structure-guided mutational analysis identifies the α2-L2 region of the HFD as a major determinant of paralog-specific partner selection. Together, our findings establish that subtle divergence within a conserved histone fold domain underlies the non-redundant integration of Taf12 paralogs into distinct coactivator complexes, revealing a mechanism by which transcriptional machinery can evolve functional specialization through gene duplication.

## Introduction

The transcription of protein-coding genes by RNA polymerase II (Pol II) is a tightly regulated process, requiring the coordinated action of general transcription factors (GTFs) and coactivators (1). Central among these is the TFIID complex, a multiprotein assembly composed of the TATA-binding protein (TBP) and 14 TBP-associated factors or TAFs. While TBP functions as a core DNA-binding component, TAFs confer functional diversity and regulatory specificity to the TFIID complex (2-4). Five of the TAFs, namely TAF5, -6, -9, -10 and TAF12 are shared between TFIID and multifunctional SAGA complex (5-7). A major structural feature shared among several TAFs is the histone-fold domain (HFD). These histone-fold TAFs (HF-TAFs) assemble into histone-like heterodimeric pairs TAF4/TAF12,TAF6/TAF9, TAF11/13, TAF8/TAF10 and TAF3/TAF10 that form the architectural scaffold of TFIID and contribute to its modular assembly (2-4). Similarly, in SAGA complex, TAFs assemble into distinct specific heterodimeric pairs through Histone fold domain interactions: TAF6/TAF9, Ada1/TAF12 and Spt7/TAF10 that form the architectural scaffold of the complex (5, 7-10). Despite sharing the five HF-TAFs, TFIID and SAGA architectural scaffold form distinct structures mostly due to the incorporation of alternative interaction partners (6, 7) Intriguingly, many TAFs exist as paralogs or are expressed in multiple isoforms, which differ in sequence, structure, and tissue distribution leading to formation of alternate TFIID-like complexes and modulating transcription in a cell-type-specific, developmental stage-specific, or signal-dependent manner (11-13). Moreover, some paralogous TAFs also show complex specific specialization with either part of TFIID or SAGA complex (14, 15).

*Candida albicans* is a critical priority group human fungal pathogen (16) that lives asymptomatically in healthy humans but upon immunocompromisation can lead to life-threatening infections such as Candidiasis and Candidemia (17). Unlike most fungi and metazoans (excluding Plants), *C. albicans* genome encode two TAF12 paralogs TAF12 and TAF12L with distinct biological functions (6, 18-21). Taf12 is essential for viability, whereas Taf12L is dispensable under standard growth conditions but required for oxidative stress tolerance (19). Gene duplication followed by subfunctionalization or neofunctionalization is a major driver of regulatory evolution, particularly in transcriptional networks (22, 23). For pathogens such as *C. albicans*, which must rapidly remodel gene expression programs in response to fluctuating host environments, diversification of core transcriptional machinery may provide a selective advantage (24).

Although previous work established preferential association of Taf12 with TFIID and Taf12L with SAGA (19), it is unclear whether this specificity reflects non-redundant association. Moreover, the molecular determinants that enforce paralog-specific incorporation into TFIID versus SAGA are unknown. Given the multitude of transcriptional complexes wherein histone fold domains mediate subunits interactions and role of these complexes in transcription, it is crucial to understand how such specificity could be mediated by the conserved Histone fold domains (25, 26).

Here, we investigate the evolutionary and structural basis of Taf12 paralog specialization in *C. albicans*. Using conditional depletion, ectopic expression, and co-immunoprecipitation, we show that Taf12 and Taf12L are non-redundant for TFIID and SAGA incorporation, respectively, even in the absence of the alternate paralog. We demonstrate that the conserved histone fold domains of each paralog are sufficient for biological function and complex incorporation yet exhibit intrinsic differences in interaction specificity with Taf4 and Ada1. Through structure-guided mutational and chimera analyses, we identify the α2-L2 region of the histone fold domain as a critical determinant of paralog-specific heterodimerization. Together, our findings reveal how modest sequence divergence within a conserved architectural motif can drive functional specialization of transcriptional coactivators, providing insight into the evolution of transcriptional regulation and its adaptation to pathogenic lifestyles.

## Results

### *CaTAF12* variants are non-redundant for their association with TFIID and SAGA complex

The two *TAF12* variants of *C. albicans; TAF12* and *TAF12L* show a functionally specialized association with TFIID and SAGA complex respectively (19). To uncover if this association is strictly non-redundant, we examined the biochemical association of each CaTaf12 variant respective to TFIID and SAGA complex in the absence of other. Specifically, we assessed Taf12’s ability to associate with TFIID and SAGA upon *TAF12L* depletion, and vice versa.

To this end, we generated strains in which either Taf12 or Taf12L was expressed from a maltose-regulatable promoter (*P*_*MAL2*_), while TFIID and SAGA specific subunits, Taf11 and Ada2 respectively, were TAP-tagged. The resulting strains; P_*MAL2*_*-TAF12L* Ada2-TAP, P_*MAL2*_*-TAF12* Ada2-TAP, P_*MAL2*_*-TAF12L* Taf11-TAP and P_*MAL2*_*-TAF12* Taf11-TAP as well as the control strains Ada2-TAP (in WT) and Taf11-TAP (in WT) exhibited comparable levels of Ada2-TAP and Taf11-TAP expression **(Figure S1 A-C)**. Next we grew these strains along with control strains WT (untagged), P_*MAL2*_*-TAF12L*, P_*MAL2*_*-TAF12*, in YP media with glucose, thereby depleting either Taf12L or Taf12 and then TAP-pulled down either Ada2 or Taf11 followed by TEV cleavage. Inputs and immunoprecipitations confirmed successful enrichment of the TAP-tagged proteins and their TEV-cleaved counterparts respectively **(Figure 1 A-B; lanes 4-6 and 10-12)**.

**Figure 1.**
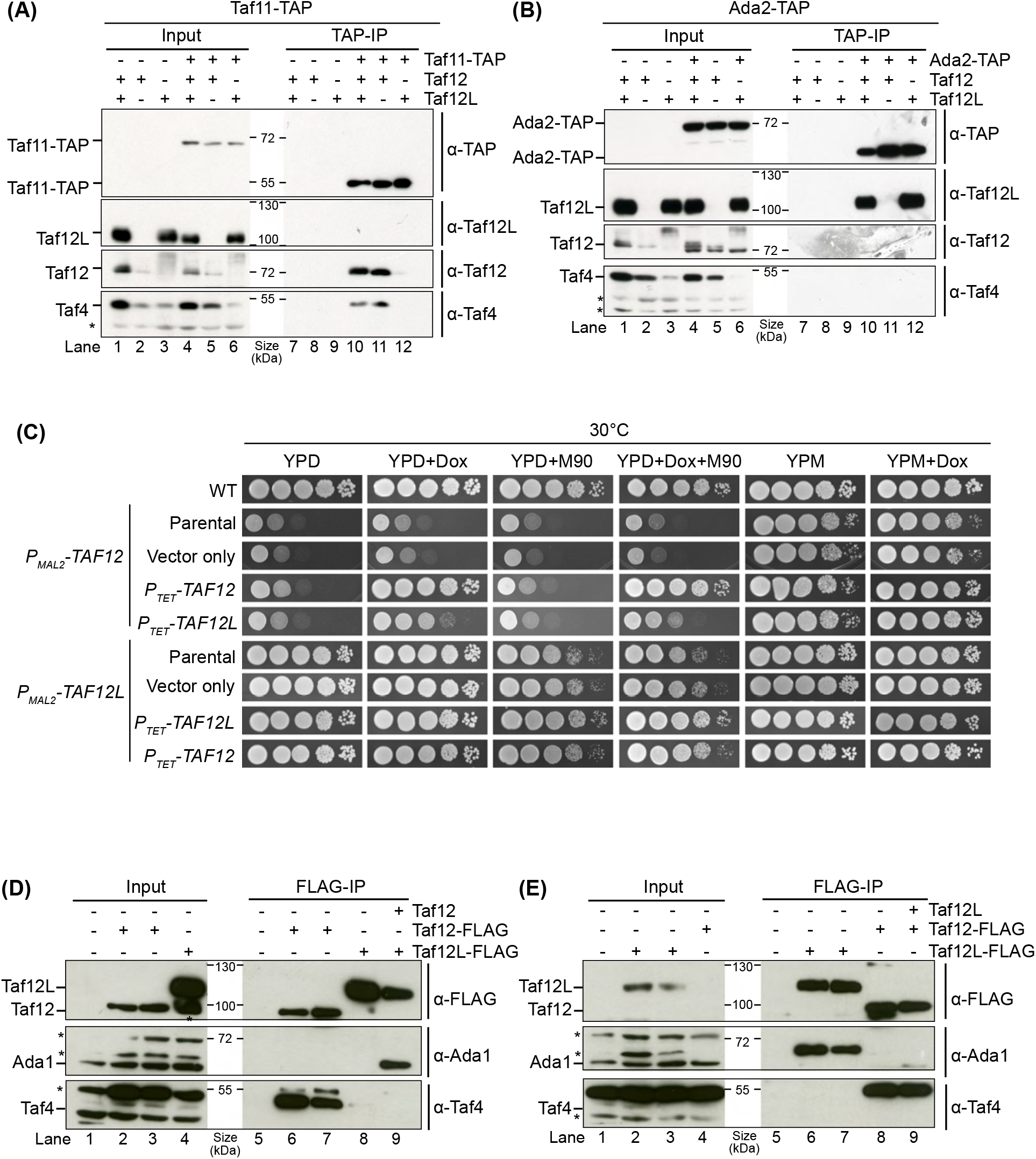
*CaTAF12* variants are non-redundant for their association with TFIID and SAGA complex. (A) Co-immunoprecipitation analysis of Taf12L association with Taf11 in absence of Taf12. 5mg equivalent of Whole cell extracts prepared from 7h YPD grown strains WT untag (SN95), P_*MAL2*_-*TAF12L* (ISC11), P_*MAL2*_-*TAF12* (ISC12), Taf11-TAP in SN95 (SDC28), Taf11-TAP in ISC11 (SDC30) and Taf11-TAP in ISC12 (SDC34) were immunoprecipitated with IgG-Sepharose beads and eluted by incubating overnight with AcTEV protease for TEV-cleavage. 50µg equivalent protein extract from each strain was used as input while 20% of the total IP elute was used. The samples were resolved on 10% SDS-PAGE gel and membranes were probed with anti-Taf12L (1:3000), anti-TAP (1:3000), anti-Taf12 (1:3000) and anti-Taf4 (1:3000). Lane 1-6 input: SN95, ISC11, ISC12, SDC28, SDC30, SDC34 input respectively. Lane 7-12 IP: SN95, ISC11, ISC12, SDC28, SDC30 and SDC34 respectively. (B) Co-immunoprecipitation analysis of Taf12 association with Ada2 in absence of Taf12L. 5mg equivalent of Whole cell extracts prepared from 7h YPD grown strains WT untag (SN95), P_*MAL2*_-*TAF12L* (ISC11), P_*MAL2*_-*TAF12* (ISC12), Ada2-TAP in SN95 (SDC27), Ada2-TAP in ISC11 (SDC29) and Ada2-TAP in ISC12 (SDC33) were immunoprecipitated with IgG-Sepharose beads, resolved on 10% SDS-PAGE gel and membranes probed with different antibodies as in (A). Lane 1-6 input: SN95, ISC11, ISC12, SDC27, SDC29 and SDC33 respectively. Lane 7-12 IP: SN95, ISC11, ISC12, SDC27, SDC29 and SDC33 respectively. (C) Spot assay analysis of growth of *P*_*TET*_ expressed Taf12 and Taf12L in Taf12 variants shut-off mutants. Strains WT (SN95), P_*MAL2*_*-TAF12* (ISC12), P_*TET*_*-TAF12* in ISC12 (SDC16), P_*TET*_*-TAF12L* in ISC12 (SDC15), vector only control (SDC6), P_*MAL2*_*-TAF12L* (ISC11), P_*TET*_*-TAF12L* in ISC11 (SDC13), P_*TET*_*-TAF12* in ISC11 (SDC14) and vector only control (SDC3) were grown in YPM liquid media till saturation and then serial dilutions were spotted on YPD, YPD+Doxycycline (50µg/ml), YPD+menadione (90µM), YPD+Doxycycline+menadione, YPM and YPM+Doxycycline plates. The plates were incubated at 30°C and imaged. (D) Co-immunoprecipitation analysis of *P*_*TET*_ expressed Taf12L and Taf12 in absence of Taf12. 5mg equivalent of Whole cell protein extracts prepared from 7h YPD+Doxycycline grown strains *TAF12-*FLAG_3_ (SKC6), *TAF12L-*FLAG_3_ (SKC3), P_*MAL2*_*-TAF12* (ISC12), P_*TET*_-*TAF12L-*FLAG_3_ in ISC12 (SDC15) and P_*TET*_-*TAF12-*FLAG_3_ in ISC12 (SDC16) were immunoprecipitated with FLAG-M2 affinity gel and eluted using 3XFLAG peptide. 100µg equivalent protein extract from each strain was used as input while 15% of the total IP elute was used. Membrane was probed with anti-FLAG (1:1000), anti-Ada1 (1:1000) and anti-Taf4 (1:3000). Lane 1-4 Input: ISC12, SKC6, SDC16, SDC15 respectively. Lane 5-9 IP: ISC12, SKC6, SDC16, SDC15 and SKC3 respectively. (E) Co-immunoprecipitation analysis of *P*_*TET*_ expressed Taf12L and Taf12 in absence of Taf12L. 5mg equivalent of Whole cell protein extracts prepared from 7h YPD+Doxycycline grown strains *TAF12-*FLAG_3_ (SKC6), *TAF12L-*FLAG_3_ (SKC3), P_*MAL2*_-*TAF12L* (ISC11), P_*TET*_-*TAF12L-*FLAG_3_ in ISC11 (SDC13) and P_*TET*_-*TAF12-*FLAG_3_ in ISC11(SDC14) were immunoprecipitated with FLAG-M2 affinity gel and eluted using 3XFLAG peptide. 100µg equivalent protein extract from each strain was used as input while 50% of the total IP elute was used. Membrane was probed with anti-FLAG (1:1000), anti-Ada1 (1:1000) and anti-Taf4 (1:3000). Lane 1-4 Input: ISC11, SKC3, SDC13, SDC14 respectively. Lane 5-9 IP: ISC11, SKC3, SDC13, SDC14 and SKC6 respectively.

We found that while Taf12 efficiently co-immunoprecipitated with Taf11-TAP irrespective of presence or absence of Taf12L **(Figure 1A; cf. lane 10 vs 11)**, Taf12L did not immunoprecipitate with Taf11-TAP even in the absence of Taf12 **(Figure 1A; cf. lanes 10 and 12)**. Moreover, Taf4 a known TFIID component, also failed to co-immunoprecipitate with Taf11-TAP in the absence of Taf12, suggesting that Taf12-Taf4 heterodimerization is essential for Taf4 incorporation into TFIID **(Figure 1A; lane 12)**. Similarly, Ada2-TAP efficiently pulled down Taf12L regardless of presence and absence of Taf12 **(Figure 1B; cf. lane 10 vs 12)**, but did not associate with Taf12 even when Taf12L was depleted **(Figure 1B; cf. lanes 10-11)**. Additionally, Taf4 was not pulled down by Ada2-TAP in any condition **(Figure 1A; lanes 10-12)** confirming that Taf4 is not part of the *Ca*SAGA complex. These results thus established that in vivo the two *C. albicans* Taf12 variants are non-redundant for their association with TFIID and SAGA complex.

To further investigate the functional specificity of Taf12 variants, we ectopically expressed 3X-FLAG tagged Taf12 and Taf12L from a doxycycline inducible *TET* promoter in Taf12 or Taf12L depleted strains **(Figure S1D i-iv)**. Growth assays and co-immunoprecipitation experiments were performed to assess functional rescue and complex incorporation. Growth assay on doxycycline containing YPD plates revealed that ectopically expressed Taf12-FLAG from *TET* promoter fully complemented the poor growth phenotype of Taf12 depletion **(Figure 1C; row 4; YPD+Dox)**. Similarly, ectopically expressed Taf12L-FLAG restored growth under oxidative stress caused by Taf12L depletion **(Figure 1C; row 8; YPD+Dox+menadione)**.

Interestingly, ectopic expression of Taf12L-FLAG partially rescued the Taf12 depletion phenotype **(Figure 1C; cf. row 5 vs 3; YPD+Dox)**, while Taf12-FLAG failed to rescue the oxidative stress phenotype of Taf12L-depleted cells **(Figure 1C; cf. row 9 vs 7; YPD+Dox+M90)**. This asymmetric complementation might result from poor expression of Taf12-FLAG from the TET promoter in Taf12L-depleted strains, while Taf12L-FLAG was expressed at higher levels in Taf12-depleted cells **(Figure S1E-F)**. This trend persisted in *taf12LΔ/Δ* mutant backgrounds, where both Taf12L and Taf12 showed poor *TET*-driven expression **(Figure S2A)**. Nonetheless, Taf12L-FLAG could fully rescue poor growth of *taf12lM/M* mutant’s while Taf12-FLAG did not show any effect on poor growth **(Figure S2B; cf. row 3 vs 4-5 YPD+Dox)**. To rule out promoter-specific effects, we used the methionine- and cysteine-repressible *P*_*MET3*_ promoter to express Taf12-FLAG and Taf12L-FLAG in *taf12LΔ/Δ* mutants. We observed better expression of Taf12L-FLAG from *P*_*MET3*_ but not for Taf12-FLAG **(Figure S2C; lanes 9-10 vs 11-12)**. Again we saw only Taf12L-FLAG but not Taf12-FLAG fully complemented *taf12lM/M* mutant’s poor growth **(Figure S2D; cf. row 3 vs 4)**.

We next tested if the partial complementation of Taf12 depletion by ectopic expression of Taf12L-FLAG is due to incorporation of Taf12L-FLAG into TFIID complex. To this end, we did FLAG co-immunoprecipitation of Taf12L-FLAG and Taf12-FLAG expressing strains under Taf12 depletion conditions (that is in presence of Glucose and doxycycline). Western blot after co-IP revealed that under Taf12 depletion revealed that Taf12-FLAG, like endogenous Taf12, specifically pulled down Taf4 but not Ada1 **(Figure 1D; cf. lanes 6 and 7; Taf4 vs Ada1 blot)** indicating specific interaction with Taf4 and incorporation into TFIID. Interestingly Taf12L-FLAG expressed from *TET* promoter also pulled down Taf4, albeit less efficiently than Taf12-FLAG suggesting partial incorporation into TFIID **(Figure 1D and S2E; cf. lane 8 vs 7; Taf4 blot)**. Moreover, Taf12L-FLAG also competed with endogenous Taf12L for Ada1 binding, showing incorporation into SAGA, though again to a lesser extent **(Figure S2E; cf. lanes 8 and 9; Ada1 blot)**.

Similarly, we tested if the lack of suppression of Taf12L depletion by ectopic expression of Taf12-FLAG is due to non-incorporation of Taf12-FLAG into SAGA complex. Therefore, we did FLAG co-immunoprecipitation of Taf12L-FLAG and Taf12-FLAG expressing strains under Taf12L depletion conditions (in presence of Glucose and doxycycline). Western blot after co-IP revealed that both endogenous Taf12L-FLAG, as well as *P*_*TET*_ expressed Taf12L-FLAG can very specifically pull down only Ada1 but not Taf4 **(Figure 1E; cf. lanes 6 and 7; Taf4 vs Ada1 blot)** indicating specific interaction with Ada1 and incorporation in SAGA complex. However, *P*_*TET*_ expressed Taf12-FLAG did not pull down Ada1 but pulled down Taf4 similar to endogenous Taf12-FLAG **(Figure 1E; cf. lanes 8 and 9)**. Together, these results demonstrate that *C. albicans* Taf12 and Taf12L are functionally non-redundant, each specifically required for incorporation into TFIID and SAGA complexes, respectively. Partial cross-complementation is limited and appears to be context and expression level dependent, underscoring the specialized roles of these paralogous factors.

### TAF12 paralogs show high specificity of interaction for TAF4 and Ada1 in vitro

TFIID and SAGA complex are both multi-subunit complexes with multiple inter-subunit contacts. Taf12 in both TFID and SAGA complex apart from heterodimerizing with Taf4 and Ada1 respectively make contact with the core scaffold including Taf6-Taf9 pair and Taf5 (3-5, 7-9). To understand if the specificity of the CaTaf12 variants is due to selective heterodimerization with Taf4 or Ada1, we performed in vitro GST pull down assays using GST tagged Ada1 and Taf4 proteins as bait and 3xFLAG tagged Taf12L and Taf12 proteins as prey. Our dose-dependent pull down showed that GST-Ada1 but not GST-Taf4 or GST pulled down specifically Taf12L-FLAG_3_ **(Figure 2A; compare lanes 8-10 vs 4-6)**. Our quantitation of the pull down experiment confirmed a progressive and specific increase in Taf12L-FLAG recovery with increasing amounts of GST-Ada1 **(Figure 2B)**. Similarly, increasing amount of GST-Taf4 but not GST-Ada1 or GST pulled down specifically Taf12-FLAG with corresponding increase in amount being pulled down **(Figure 2C; lanes compare lanes 8-10 vs 4-6 and Figure 2D)**. Thus these data revealed that both Taf12 variants very specifically interact with either Ada1 or Taf4.

**Figure 2.**
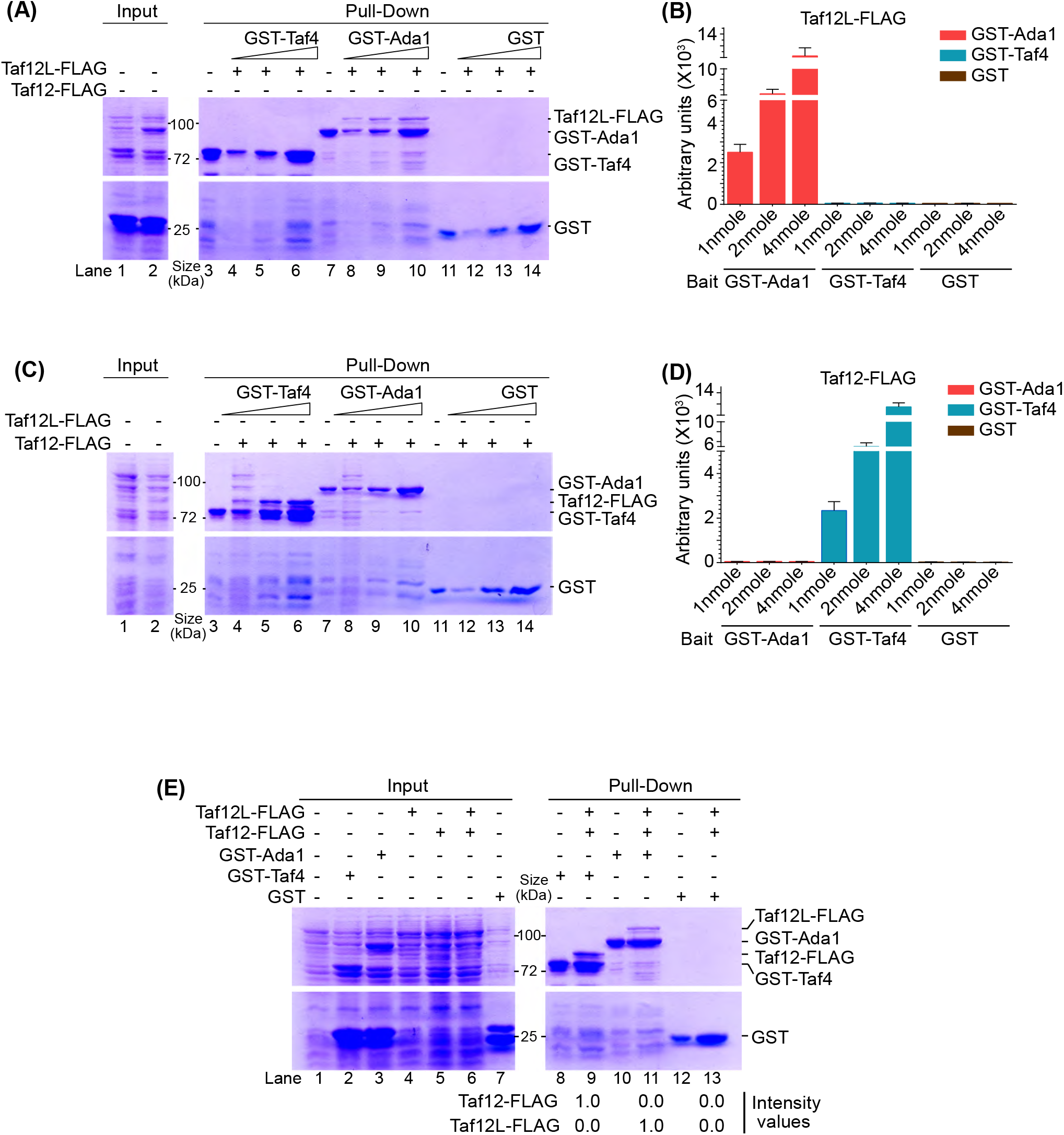
Taf12 variants show specific interaction with Taf4 and Ada1 proteins in vitro. (A) GST-Ada1 specifically pull-down Taf12L-FLAG_3_ in dose dependent manner. Equal volume of Glutathione Sepharose 4B beads was incubated with 1nmole, 2nmole and 4nmole of GST-Taf4 (Lanes 4-6), GST-Ada1 (Lanes 8-10) and GST (Lanes 12-14). Beads were washed and incubated for 2hr with 1nmole of Taf12L-FLAG_3_. Beads were boiled and run on gel. Lanes 1 and 2: input for GST-Taf4 and GST-Ada1 respectively. Lanes 3, 7 and 11: Control Pull down of GST-Taf4, GST-Ada1 and GST respectively. (B) Quantification of amount of Taf12L-FLAG_3_ being pulled down with different amounts of GST-Ada1, GST-Taf4 and GST respectively (from A). The units on y-axis represent the ImageJ intensity values. (C) GST-Taf4 specifically pull-down Taf12-FLAG_3_ in dose dependent manner. Equal volume of Glutathione Sepharose 4B was incubated with 1nmole, 2nmole and 4nmole of GST-Taf4 (Lanes 4-6), GST-Ada1 (Lanes 8-10) and GST (Lanes 12-14). Beads were washed and incubated with 1nmole of Taf12-FLAG_3_. Beads were boiled and run on gel. Lanes 1 and 2: input for GST-Taf4 and GST-Ada1 respectively. Lanes 3, 7 and 11: Control Pull down of GST-Taf4, GST-Ada1 and GST respectively FLAG_3_. (D) Quantification of amount of Taf12-FLAG_3_ being pulled down with different amounts of GST-Ada1, GST-Taf4 and GST respectively (from B). The units on y-axis represent the ImageJ intensity values. (E) CaTaf12L selectively interacts with Ada1 while CaTaf12 specifically interacts with Taf4. Equal volume of Glutathione Sepharose 4B beads was incubated with 1nmole of GST-Taf4 (Lanes 8 and 9), GST-Ada1 (Lanes 10 and 11) and GST (Lanes 12 and 13). Beads were washed and incubated with a mixed lysate containing 1nmole each of Taf12L-FLAG_3_ and Taf12-FLAG_3_. Beads were boiled and run on gel. Lanes 2-7: input for GST-Taf4, GST-Ada1, Taf12L-FLAG_3_, Taf12-FLAG_3_, mixed input for Taf12L-FLAG_3_ and Taf12-FLAG_3_ and GST respectively. Lane 1: uninduced control, Lanes 8, 10 and 12: Control Pull down of GST-Taf4, GST-Ada1 and GST. The values below the gel represent the ImageJ intensity values.

To further test this specificity of heterodimerization, we performed GST pull down assay using a mixed extract containing both Taf12L-FLAG and Taf12-FLAG. Notably, we observed a very specific pull down of Taf12L-FLAG with GST-Ada1 and of Taf12-FLAG with GST-Taf4 **(Figure 2E compare lane 9 and 11)**. Together, our *in vitro* interaction assays showed that the two Taf12 variants exhibit a high inherent specificity and selectivity of interaction with respect to Taf4 and Ada1 as heterodimerization partners.

### Histone Fold domains (HFDs) of Ca*TAF12* variants are sufficient for their molecular function

All Taf12 and Taf12 like proteins contain a highly conserved H2B like histone fold (HF) domain in their C-terminus (19). In higher eukaryotes such as human, *Drosophila* etc. Taf12 is a small protein composed almost entirely of the HF domain. On the other hand, *S. cerevisiae TAF12* possess a longer N-terminus apart from the HF domain that have interaction motifs for Tra1 (7, 9). However, previously it has been shown that C-terminal conserved region of *yTAF12* containing the histone fold (HF) domain is sufficient for complementation of *taf12*Δ lethality (19). Both *CaTAF12* variants possess the similar N-terminus as of *yTAF12* along with the HF domain. While the HF domain is highly conserved, the N-terminus of *CaTAF12* variants possess significant sequence differences (19). Therefore, we wanted to assess the functional significance of the conserved HF domains and the non-conserved sequences of the two *TAF12* variants in *C. albicans*. To this end, expressed only 3XFLAG tagged HF domains of one or the other *TAF12* variant from *TET* promoter in the background of *TAF12* or *TAF12L* shut-off mutant strains similar to that for full length versions **(Figure S2A)**.We observed a robust expression of both HFD-*TAF12-*FLAG and HFD-*TAF12L-*FLAG, under *TAF12* depletion conditions but very poor expression under *TAF12L* depletion conditions **(Figure S3)**. Importantly, despite the poor expression, HFD-*TAF12L* fully complemented menadione induced poor growth phenotype of *TAF12L* depletion **(Figure 3A compare rows 7 vs 6 YPD+Dox+menadione)**. Similarly, HFD-*TAF12* fully complemented poor growth phenotype of *TAF12* depletion **(Figure 3A compare rows 3 vs 2 YPD+Dox)**. Moreover, ectopic expression of *HFD-TAF12L* partially rescued the *TAF12* depletion phenotype, mimicking full-length full length *TAF12L* **(Figure 3A; row 4, YPD+Dox)**. In contrast, *HFD-TAF12*, like full-length *TAF12*, did not rescue the *TAF12L* depletion phenotype in the presence of oxidative stress **(Figure 3A; row 8; YPD+Dox+menadione)**. These results established that the Histone fold (HF) domains of *CaTAF12* variants are sufficient to mediate their biological functions.

**Figure 3.**
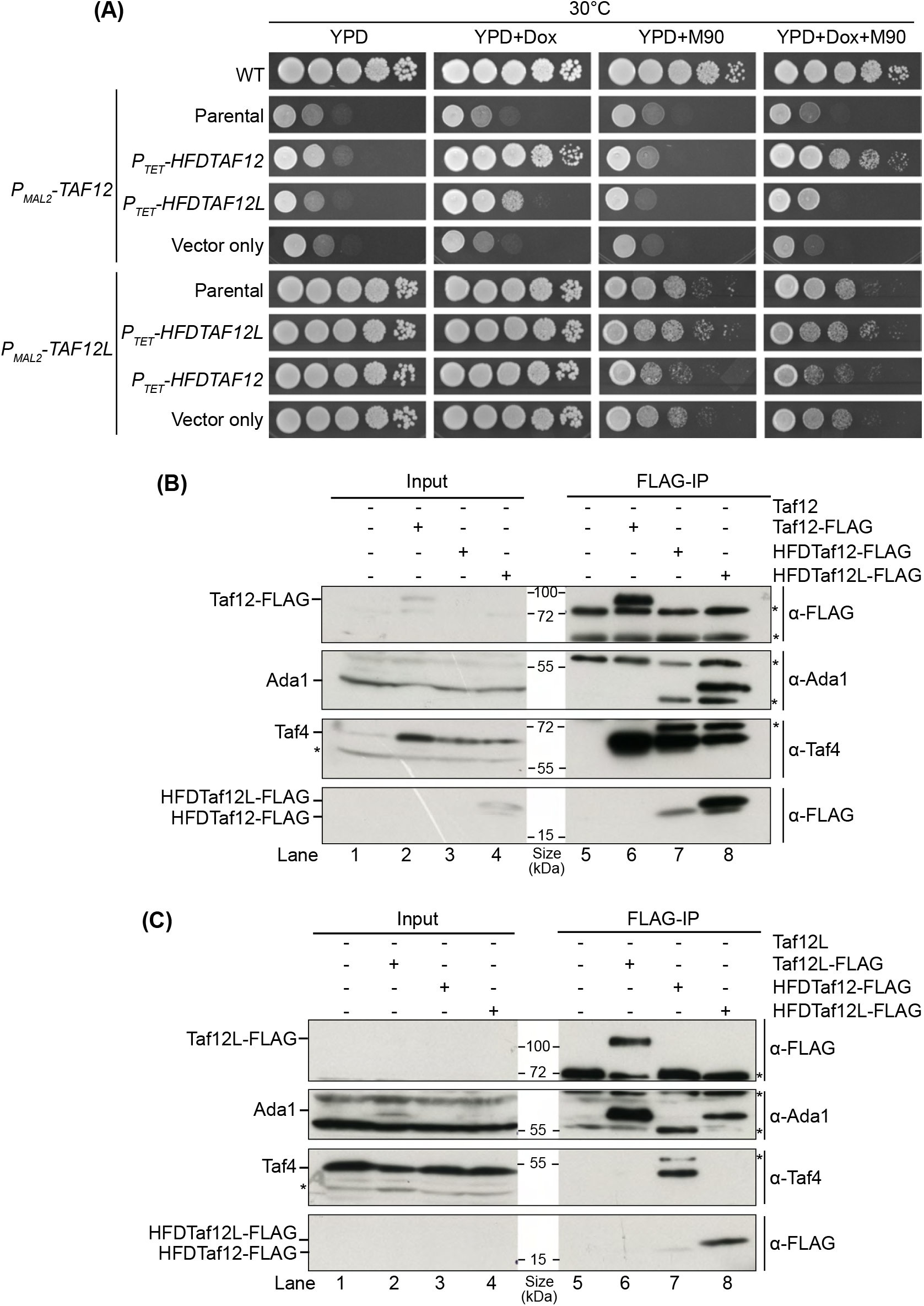
Histone Fold domains (HFDs) of *CaTAF12* variants are sufficient for their biochemical association and molecular function. (A) *P*_*TET*_ expressed HFDs can complement absence of respective *TAF12* variants shut-off mutants. Effect of HFD-*TAF12* and HFD-*TAF12L* complementation on growth of *TAF12* shut-off mutant. Strains WT (SN95), P_*MAL2*_*-TAF12* (ISC12), P_*TET*_*-*HFD*TAF12* in ISC12 (SDC5), P_*TET*_*-*HFD*TAF12L* in ISC12 (SDC8), vector only control (SDC6), P_*MAL2*_*-TAF12L* (ISC11), P_*TET*_*-*HFD*TAF12L* in ISC11 (SDC7), P_*TET*_*-*HFD*TAF12* in ISC11 (SDC2) and vector only control (SDC3) were grown in YPM liquid media till saturation and then serial dilutions were spotted on YPD, YPD+Doxycycline (50µg/ml), YPD+menadione (90µM), YPD+Doxycycline+menadione plates. The plates were incubated at 30°C and imaged. (B) Co-immunoprecipitation analysis of *P*_*TET*_ expressed HFDs of *TAF12* variants in absence of Taf12. 5mg equivalent of Whole cell protein extracts prepared from 7h YPD+Doxycycline grown strains *TAF12-*FLAG_3_ (SKC6), P_*MAL2*_-*TAF12* (ISC12), P_*TET*_-HFD*TAF12-*FLAG_3_ in ISC12 (SDC5) and P_*TET*_-HFD*TAF12L-*FLAG_3_ in ISC12 (SDC8) were immunoprecipitated with FLAG-M2 affinity gel and eluted using 3XFLAG peptide. 150µg equivalent protein extract from each strain was used as input while 50% of the total IP elute was resolved on 4-15% gradient SDS-PAGE gels. Membranes were probed with anti-FLAG (1:1000), anti-Taf4 (1:3000) and anti-Ada1 (1:1000). Lane 1-4 input: ISC12, SKC6, SDC5 and SDC8 respectively. Lane 5-8 IP: ISC12, SKC6, SDC5 and SDC8 respectively. (C) Co-immunoprecipitation analysis of *P*_*TET*_ expressed HFDs of *TAF12* variants in absence of Taf12L. 5mg equivalent of Whole cell protein extracts prepared from 7h YPD+Doxycycline grown strains *TAF12L-*FLAG_3_ (SKC3), P_*MAL2*_-*TAF12L* (ISC11), P_*TET*_-HFD*TAF12-*FLAG_3_ in ISC11 (SDC2) and P_*TET*_-HFD*TAF12L-*FLAG_3_ in ISC11 (SDC7) were immunoprecipitated with FLAG-M2 affinity gel and eluted using 3XFLAG peptide. 150µg equivalent protein extract from each strain was used as input while 50% of the total IP elute was resolved on 4-15% gradient SDS-PAGE gels. Membranes were probed with anti-FLAG (1:1000), anti-Taf4 (1:3000) and anti-Ada1 (1:1000). Lane 1-4 input: ISC11, SKC3, SDC2 and SDC7 respectively. Lane 5-8 IP: ISC11, SKC3, SDC2 and SDC7 respectively.

To test whether the HF domains are sufficient for incorporation into the TFIID and SAGA complexes we performed FLAG co-immunoprecipitation of *P*_*TET*_-driven HFD-Taf12 and HFD-Taf12L following depletion of the corresponding endogenous proteins. We observed that HFD-Taf12 specifically immunoprecipitated Taf4 similar to the endogenous Taf12 **(Figure 3B; compare lanes 6 and 7)** but not Ada1. Importantly, we observed that P_*TET*_ expressed HFD-Taf12L also immunoprecipitated Taf4, though at a lower level than HFD-Taf12 **(Figure 3B; compare lanes 8 vs 7; Taf4 blot)**. Moreover, in the same immunoprecipitation HFD-Taf12L also pulled down Ada1. Correspondingly, we observed HFD-Taf12L specifically immunoprecipitated Ada1 similar to the endogenous Taf12 **(Figure 3C; compare lanes 6 and 8; Ada1 blot)**. Interestingly, while HFD-Taf12L could interact with Taf4 in the absence of Taf12, this interaction was not observed when Taf12 was present **(Figure 3C vs 3B; compare lane 8)**. On the contrary, P_*TET*_ expressed HFD-Taf12 immunoprecipitated Taf4, but not Ada1 **(Figure 3B; compare lanes 8 vs 7)**. Together, these results established that the histone fold (HF) domains of CaTaf12 variants are sufficient for the interaction with Taf4 and Ada1 and incorporation into the TFIID and SAGA complex respectively.

### Histone fold (HF) domain of Taf12 can interact both Taf4 and Ada1 proteins *in vitro*

The earlier studies on different *TAF* heterodimers (27-33) have shown that TAFs heterodimerize though their histone fold domains or HFDs. Additionally, having established that HFD-Taf12L can interact with Taf4, we wanted to look for the sufficiency of HFDs in the heterodimerization and specificity of Ca*TAF12* variants. Therefore, we did in vitro GST-pull down assays using GST tagged Histone fold (HFs) of Taf4 and Ada1 as bait proteins and purified His_6_ tagged HF domains of Taf12L and Taf12 as prey. We found that only HFD-Ada1 but not HFD-Taf4 or GST specifically pulled down HFD-Taf12L **(Figure 4A; compare lanes 9-11 vs 5-7 and 14-16)**. Quantitation of amount of HFD-Taf12L pulled down showed that with increasing amount of HFD-Ada1, increasing amount of HFD-Taf12L pulled down **(Figure 4B)**. Interestingly, we observed that both HFD-Taf4 and HFD-Ada1 pulled down HFD-Taf12 **(Figure 4C; compare lanes 9-11 vs 5-7 and 14-16)**. Furthermore, quantitation of pull down experiment showed that increased amount of both HFD-Taf4 and HFD-Ada1 correspondingly pulled down increased amount of HFD-Taf12 **(Figure 4D)**.

**Figure 4.**
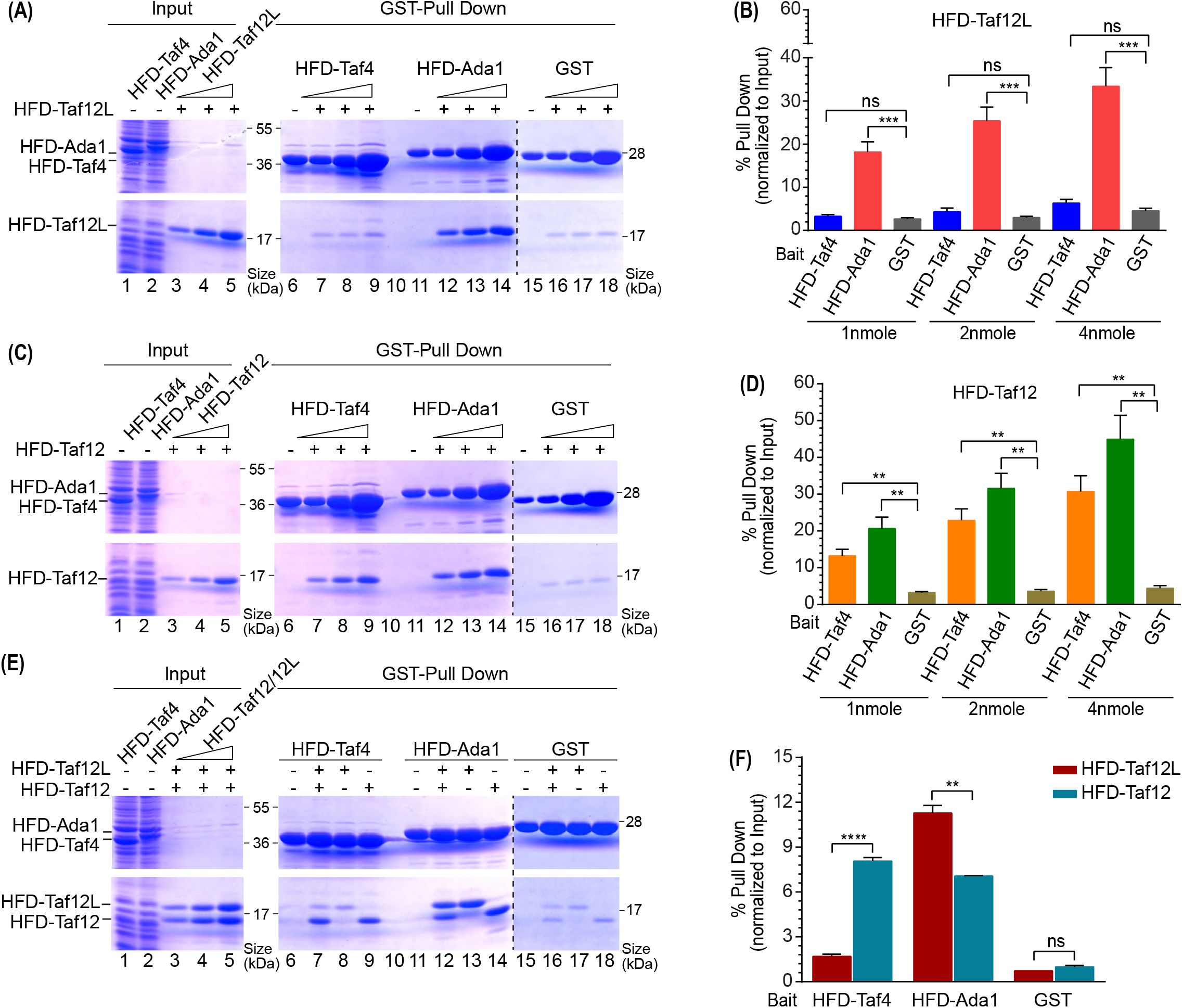
Histone fold domain of Taf12 can interact with histone fold domain of both Taf4 and Ada1 in vitro. **(A)** HFDTaf12L can be pulled down specifically by HFDAda1. Equal volume of Glutathione Sepharose 4B beads was incubated with 1nmole, 2nmole and 4nmole of GST-HFDTaf4 (Lanes 7-9), GST-HFDAda1 (Lanes 12-14) and GST (Lanes 16-18). Beads were washed and incubated with 1nmole of purified HFDTaf12L. Beads were boiled and resolved on 4-15% SDS-PAGE gel. Lane 1: input for GST-Taf4, Lane 2: input for GST-Ada1, Lanes 3-5: three dilutions of input for HFDTaf12L-His_6_, lanes 4, 8 and 15 represents GST-HFDTaf4, GST-HFDAda1 and GST bound beads respectively. **(B)** Quantitation of amount of HFDTaf12L from (A) being pulled down with different amounts of GST-HFDAda1, GST-HFDTaf4 and GST. The y-axis represents the percentage amount of HFDTaf12L pulled down normalized to the amount used as input. **(C)** HFDTaf12 can be pulled down both by HFDTaf4 and HFDAda1. Equal volume of Glutathione Sepharose 4B beads was incubated with 1nmole, 2nmole and 4nmole of GST-HFDTaf4 (Lanes 7-9), GST-HFDAda1 (Lanes 12-14) and GST (Lanes 16-18). Beads were washed and incubated with 1nmole of purified HFDTaf12L. Beads were boiled and resolved on 4-15% SDS-PAGE gel. Lane 2: input for GST-Ada1, Lanes 3-5: three dilutions of input for HFDTaf12-His_6_, lanes 4, 8 and 15 represents GST-HFDTaf4, GST-HFDAda1 and GST bound beads respectively. **(D)** Quantitation of amount of HFDTaf12 from (C) being pulled down with different amounts of GST-HFDAda1, GST-HFDTaf4 and GST. The y-axis represents the percentage amount of HFDTaf12L pulled down normalized to the amount used as input. **(E)** HFD-Taf12L competitively interact with HFD-Ada1. Equal volume of Glutathione Sepharose 4B beads was incubated with 1nmole of GST-HFDTaf4 (Lanes 7-9), GST-HFDAda1 (Lanes 12-14) and GST (Lanes 16-18). Beads were washed and incubated with 1nmole of purified HFDTaf12L (lanes 8, 13 and 17), 1nmole of purified HFDTaf12 (lanes 9, 14 and 18) and with a mixed sample containing 1nmole each of HFDTaf12L and HFDTaf12 (lanes 7, 12 and 16). Beads were boiled and resolved on 4-15% SDS-PAGE gel. Lane 1: input for GST-Taf4, Lane 2: input for GST-Ada1, Lanes 3-5: three dilutions of input for HFDTaf12-His_6_, lanes 6, 11 and 15 represents GST-HFDTaf4, GST-HFDAda1 and GST bound beads respectively. **(F)** Quantification of amount of HFDTaf12L and HFDTaf12 being pulled down with GST-HFDAda1, GST-HFDTaf4 and GST respectively. The y-axis represents the percentage amount of HFDTaf12L-His_6_ pulled down normalized to the amount used as input.

The promiscuity of HFD-Taf12 interaction with HFD-Taf4 and HFD-Ada1 prompted us to probe into selectivity of these interactions. Therefore, we performed another GST-pull down assay but by using 1:1 mixture of HFD-Taf12 and HFD-Taf12L as prey. Again we observed that HFD-Taf4 selectively pulled down HFD-Taf12 only, while HFD-Ada1 pulled down both HFD-Taf12L and HFD-Taf12 **(Figure 4E; compare lanes 9 and 11)**. The quantitation of the data showed that despite both HFD-Taf12L and HFD-Taf12 being pulled down with HFD-Ada1, a significantly higher amount of HFD-Taf12L was pulled down as compared to the HFD-Taf12 **(Figure 4F)**.

As we found that the full length Taf12 variants interactions are very specific but their HFD interactions somewhat promiscuous, we asked if HFDs of CaTaf12 variants maintain this promiscuity even for full length Taf4 and Ada1. So, we did GST pull-down assay using full length GST-Taf4 and GST-Ada1 as baits and His_6_ tagged HFDs of Taf12L and Taf12 as prey proteins. Interestingly, consistent with previous results, GST-Taf4 specifically pulled down HFD-Taf12 but GST-Ada1 pulled down both HFD-Taf12L and HFD-Taf12. Again from the 1:1 mixture of HFD-Taf12L and HFD-Taf12, GST-Ada1 pulled down significantly higher amounts of HFD-Taf12L than HFD-Taf12 **(Figure S4 A-E)**.

Thus, together these results demonstrate that the HF domains of CaTaf12 variants are sufficient to mediate specific interactions with Taf4 and Ada1. In vitro, HFD-Taf12 exhibits promiscuous binding to both partners, while HFD-Taf12L preferentially and selectively binds Ada1. However, the interaction of Ada1 with HFD-Taf12L was more selective than with HFD-Taf12.

### Histone fold domain of Taf12 variants have similar interaction interface with Taf4 and Ada1

Our genetic and biochemical analyses demonstrated that the two *C. albicans* Taf12 variants, Taf12 and Taf12L, exhibit high specificity in heterodimerization with Taf4 and Ada1, respectively, both in vivo and in vitro. This is in contrast to most higher eukaryotes and yeast species, which typically express a single Taf12 protein capable of interacting with both Taf4 and Ada1 (19, 27, 34). So, we asked what drives the specificity of interaction of the two Taf12 variants. To investigate the molecular basis of this specificity, we examined the alpha-fold generated models of Histone fold (HF) domains of two CaTaf12 variants (Taf12 and Taf12L), Taf4 and Ada1 as well as alpha-fold model of heterodimers of Histone fold (HF) domains of Taf12-Taf4, Taf12L-Taf4, Taf12-Ada1 and Taf12L-Ada1.

While the HF domain is highly conserved, only Taf12L contains characteristic polyglutamine-rich stretches and Tra1 interaction regions (TIRs) in its N-terminal region **(Figure S5A-B)**. Sequence alignment of the C-terminal HF domains from 35 Taf12 homologs across multiple species including *Candida, Saccharomyces, Schizosaccharomyces pombe, Arabidopsis thaliana, Drosophila melanogaster, Mus musculus*, and *Homo sapiens* revealed strong conservation across the three core α-helices (α1, α2, α3) and the C-terminal αC helix **(Figure S5B)**. Furthermore, many of the hydrophobic and charged residues involved in heterodimerization and salt-bridge formation (as described for hTAF12–hTAF4) (27) were conserved between the two CaTaf12 variants **(Figure S5B)**. However, we found that out of a total of 116 residues encompassing the whole conserved C-terminus, 38 residues differed in their charge property, size or aromatic nature between two CaTaf12 variants. Out of which 23 residues were conserved in three or more Taf12 proteins from *Candida species*, and are markedly different in Taf12L-like sequences **(Table S4)**.

AlphaFold models of the HFDs of Taf12 and Taf12L revealed typical histone fold topology, with three core α-helices (α1–α3) and interconnecting loops L1 and L2 **(Figure S6A i-ii)**. Moreover, superimposition of the two models revealed a very similar folding pattern for HFDs of both Taf12 and Taf12L **(Figure S6A iii)**. In contrast, the alpha-fold models of HFD of Taf4 and Ada1 showed notable differences. While Ada1 HFD showed the typical HFD conformation but with a very long loop L2 and a shorter α3 helix, HFD of Taf4 showed an atypical conformation with α3 helix assuming a different orientation extending outside **(Figure S6B i-iii)**. The structural studies has indeed shown that Taf4 has an atypical HFD with long L2 and flexible α3 helix along with an extended C-terminal helix CCTD (3, 4, 27, 34).

Modelling of the four HFD heterodimers; Taf12-Taf4, Taf12L-Taf4, Taf12-Ada1 and Taf12L-Ada1 revealed that all the four structures were plausible with high confidence and very similar in overall conformation with each assuming the canonical of HF-HF heterodimers conformations **(Figure 5)**. Notably, while in monomeric alpha model of Taf4, the α3 helix assumed an extended orientation, in both Taf12-Taf4 and Taf12L-Taf4 heterodimers this helix assumed the canonical crossover orientation **(Compare Figure 5 A i-ii vs Figure S5A iii)**. Moreover, on superimposing HFD heterodimers of Taf12-Taf4 and Taf12L-Taf4, we noted considerable difference in the orientation of α3 helix. While in Taf12-Taf4 heterodimer, this helix is

**Figure 5.**
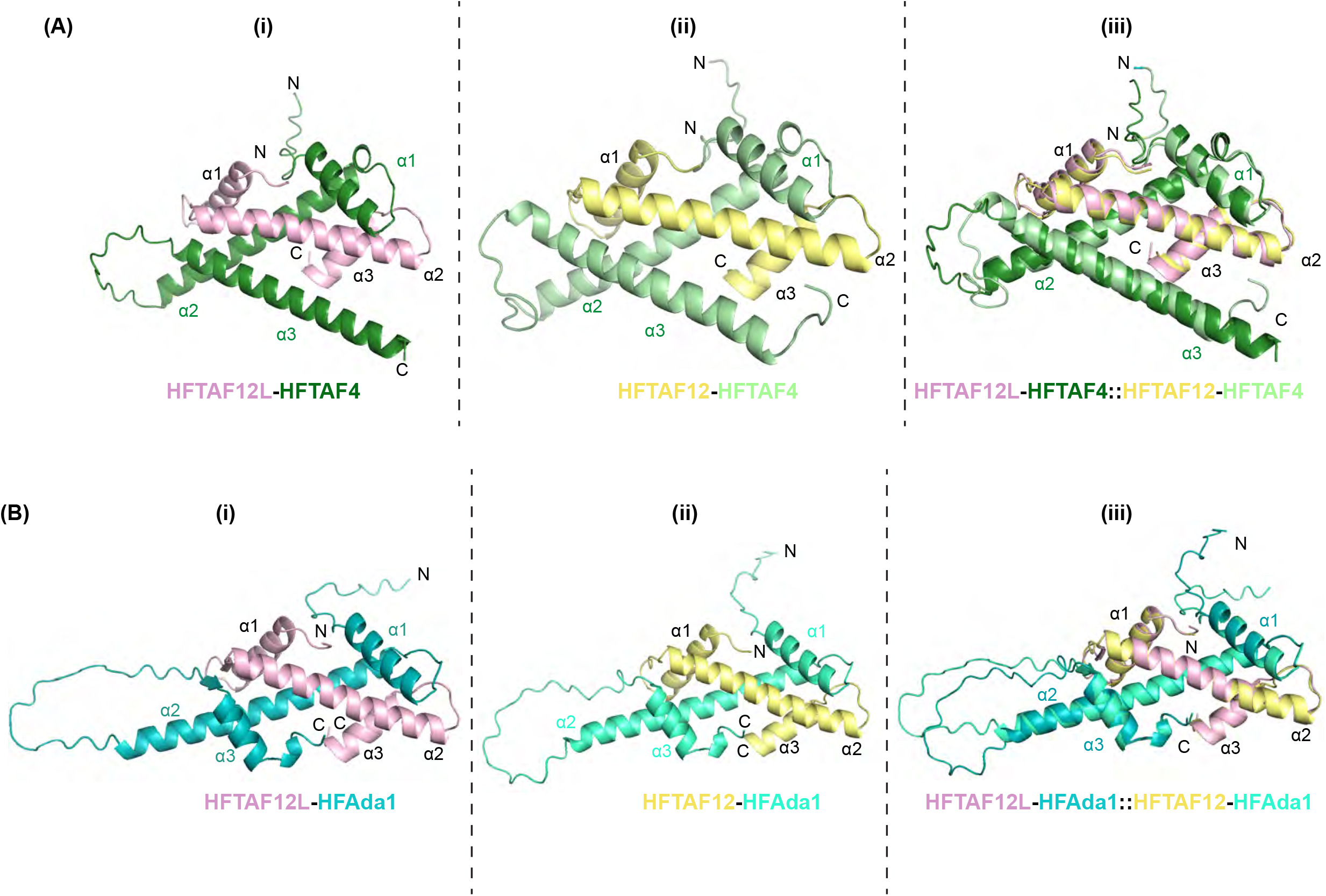
Histone fold domain of Taf12 variants have similar interaction interface with Taf4 and Ada1. **(A i-iii)** Alpha-fold model of Histone fold domains (HFD) heterodimers of *Candida albicans* Taf12 variants with HFD of Taf4. (i) heterodimer of HFD-Taf12L (in pink) and HFD-Taf4 (in green), (ii) heterodimer of HFD-Taf12 (in yellow) and HFD-Taf4 (in light green) (iii) superimposition of heterodimers from (i and ii). **(B i-iii)** Alpha-fold model of Histone fold domains (HFD) heterodimers of *Candida albicans* Taf12 variants with HFD of Ada1. (i) heterodimer of HFD-Taf12L (in pink) and HFD-Ada1 (in teal), (ii) heterodimer of HFD-Taf12 (in yellow) and HFD-Ada1 (in cyan) (iii) superimposition of heterodimers from (i and ii).

We then looked at the predicted and possible interactions between different proteins in different models at a distance of 4Ǻ. Both the Taf12 proteins showed similar type of predicted interaction interfaces with Ada1, and also with Taf4. While the interaction interfaces were different for Ada1 and Taf4 proteins, the residues involved were similar or identical for Taf12 variants. We then looked at the Taf12 variant residues that were markedly different between the two proteins as were described in previous section and the interactions that were predicted for those residues. We also mapped those residues as well as the interacting residues of Taf4 and Ada1 to the heterodimeric models. We noticed that although specific CSR residues for Taf12 variants were different in their chemical properties still they showed predicted interactions with same residue for Ada1 and similarly with same residues of Taf4 **(Table. S5)**..

Thus, our heterodimeric modelling revealed that while most of the CSR residues for Taf12 variants were different for their charge or structural properties, we were able to predict similar interactions for these residues. Only one major difference was seen for the variants but again this difference was not observed with respect to Taf4 and Ada1 models.

### α2-L2 region of HFD of Taf12 paralogs is a potential specificity determinant

While AlphaFold modelling suggested that both Taf12 and Taf12L could form stable heterodimers with either Taf4 or Ada1, our in vivo and in vitro data demonstrated distinct partner specificity. Therefore, to identify regions within the HFD responsible for functional specificity, we generated 3XFLAG tagged chimeric mutants of the two proteins **(Figure 6A i-ii)** by swapping individual helices and loop regions of two proteins. We then ectopically expressed these chimeric constructs from doxycycline inducible *P*_*TET*_ promoter in the depletion of Taf12 as previously with full length Taf12 variants and only HFD constructs **(Figure S1D)**. We observed a robust expression for all the chimeric constructs under Taf12 depletion conditions **(Figure S7)**. All Taf12-based chimeras supported robust growth under Taf12-depleted conditions, comparable to native Taf12 or HFD-Taf12 constructs **(Figure 6B compare rows 5-8 vs 3-4 YPD+doxycycline)**. In contrast, swapping the entire HFD of Taf12 with that of Taf12L only partially rescued growth, similar to the native Taf12L construct **(Figure 6B compare rows 9 vs 10 YPD+doxycycline)**.

**Figure 6.**
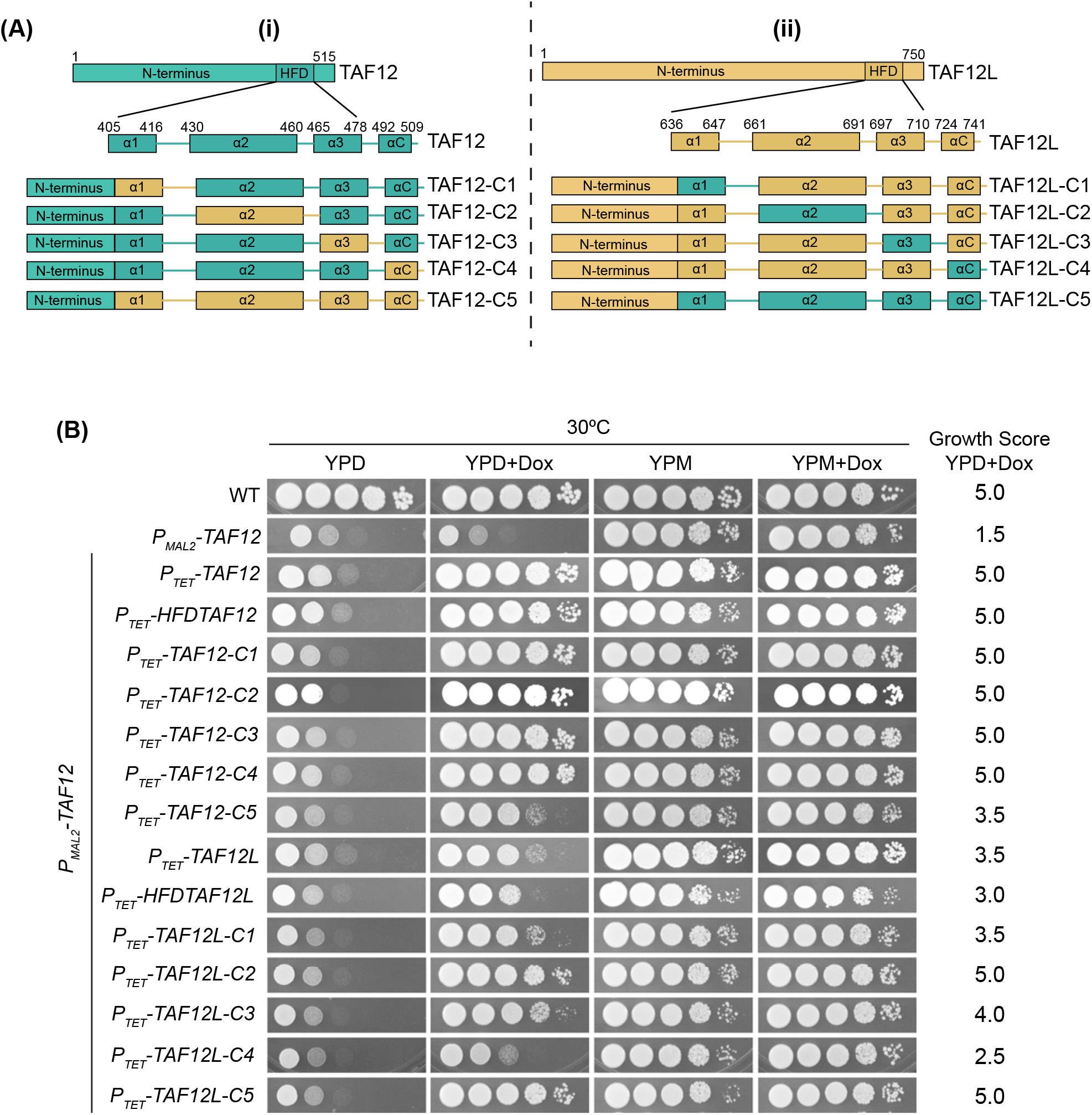
Helix swap chimera constructs of *CaTAF12* variants show differential complementation effect. **(A i-ii)** Schematic representation of CaTaf12 variants chimeric mutant constructs. (i) Chimeric constructs for Taf12. Full length Taf12 and Histone fold domain (HFD) of Taf12 is shown in teal and the regions swapped with respective Taf12L regions are shown in yellow. (ii) Chimeric constructs for Taf12L. Full length Taf12L and Histone fold domain (HFD) of Taf12L is shown in yellow and the regions swapped with respective Taf12 regions are shown in teal. (C) Spot assay analysis of growth of P_*MAL2*_*-TAF12* mutant strain expressing different chimeric constructs from (A) under *TET* promoter. All the mutant strains along with WT (SN95), and P_*MAL2*_*-TAF12* (ISC12) were grown in YPM liquid media till saturation and then serial dilutions were spotted on YPD, YPD+Doxycycline (50µg/ml), YPM and YPM+Doxycycline. The plates were incubated at 30°C and imaged. The growth was scored on a scale of 0-5 based on growth on YPD+Doxycycline plates.

A strikingly different pattern emerged with Taf12L-based chimeras. While native Taf12L only partially rescued Taf12 depletion, a Taf12L construct containing the α2–L2 region from Taf12 fully suppressed the growth defect, mimicking native Taf12 **(Figure 6B compare rows 13 vs 10 and 3 YPD+doxycycline)**. Moreover, both Taf12L construct with swapped α3-L3 or α1-L1 region partially supressed the poor growth phenotype with α3-L3 construct providing better suppression than that observed for α1-L1 construct or native Taf12L **(Figure 6B compare rows 14, 12 and 10 YPD+doxycycline)**. The Taf12L construct with swapped most C-terminal helix αC show a slightly discernible suppression of poor growth of Taf12 depletion but less than compared to native Taf12L **(Figure 6B compare rows 15 vs 10 and 2 YPD+doxycycline)**. Importantly, full replacement of the Taf12L HFD with that of Taf12 fully restored growth, equal to native Taf12 **(Figure 6B compare rows 16 vs 10 and 3 YPD+doxycycline)**. Together, these results demonstrate that while multiple regions contribute to partner specificity, the α2–L2 region within the HF domain plays a major role in defining the selective interaction of CaTaf12 variants with Taf4 and Ada1. The HF domain, particularly the α2–L2 region, is thus a key determinant of functional specificity in Taf12 heterodimerization.

## Discussion

TBP-associated factors (TAFs) are integral components of the large multisubunit co-activator complexes TFIID and SAGA. Five TAFs—TAF5, TAF6, TAF9, TAF10, and TAF12— are shared between both complexes and play a central role in defining and maintaining their core architecture. This structural integrity is mediated through TAF-TAF interactions, particularly histone fold (HF) domain–driven heterodimerization (2-4, 7, 9, 35).

### TAF12 paralog specialization in *C. albicans* coactivator complexes

Our study defines the molecular basis for the functional divergence of *Candida albicans* Taf12 paralogs; Taf12 and Taf12L in mediating their selective incorporation into the TFIID and SAGA coactivator complexes, respectively. Unlike many eukaryotes, where a single TAF12 protein is shared between both TFIID and SAGA complexes, *C. albicans* encodes two distinct TAF12 paralogs that display complex-specific associations and functions (36). The _12_ dispensability of Taf12L under standard growth conditions offers a unique opportunity to explore the mechanisms underlying paralog-specific integration into transcriptional coactivator complexes.

Our co-immunoprecipitation data show that Taf12 and Taf12L associate specifically with TFIID through Taf4 and SAGA through Ada1, respectively, and do not substitute for one another in complex formation, even in the absence of their endogenous paralog. Furthermore, overexpression of Taf12L only partially rescued the growth defects of Taf12-depleted cells and showed minimal interaction with Taf4 **(Figure 1)**. GST pull-down assays confirmed this specificity, with Taf12 exclusively binding Taf4, and Taf12L binding only to Ada1 **(Figure 2)**. These observations underscore the intrinsic selectivity in their protein–protein interactions and suggest that the paralogs function non-redundantly in vivo.

### Histone Fold Domains dictate specificity and function of *C. albicans* Taf12 paralogs

The histone fold domain (HFD) is a well-established structural motif mediating TAF-TAF interactions (3, 4, 7, 9). Our results demonstrate that the HFDs of *C. albicans* Taf12 and Taf12L are both necessary and sufficient for complex association and biological function. Expression of the isolated HFDs rescued the growth defects of their respective full-length paralog depletion and preserved interaction specificity with Taf4 or Ada1 **(Figure 3)**. This reinforces the central role of the HFD in defining complex-specific incorporation and essential function, in line with observations in budding yeast (37).

Despite overall conservation, our in vitro binding assays revealed that the two HFDs differ in interaction affinity and specificity. While Taf12L interacts robustly and exclusively with Ada1, Taf12 shows weaker, more promiscuous binding to both Ada1 and Taf4 **(Figure 4)**. Importantly, Taf12L could partially compete for Taf4 binding and rescue the growth phenotype of Taf12 depletion when overexpressed. These findings suggest that while conserved structural features allow dimerization with both partners, in vivo complex incorporation is likely governed by relative affinity and availability, potentially modulated by co-translational assembly (38).

### Structural determinants of Paralog-Specific dimerization

To identify the regions conferring specificity within the HFD, we employed structure-guided modeling and functional analysis of chimeric constructs. AlphaFold predictions supported the feasibility of all four heterodimeric combinations; Taf12-Taf4, Taf12-Ada1, Taf12L-Taf4, Taf12L-Ada1 **(Figure 5)**, suggesting that in vivo specificity may arise from subtle differences in helix or loop orientation. Functional dissection using chimeras revealed that swapping the full HFD of Taf12 with that of Taf12L (Taf12C5) resulted in partial rescue of the growth phenotype **(Figure 6)**, similar to that observed with full-length Taf12L suggesting that sequences outside the HFD play a minimal role in Taf12-specific function and complex association.

Notably replacing the α2-L2 region of Taf12L with that of Taf12 (Taf12LC2) was sufficient to fully rescue the Taf12 depletion phenotype, pointing to α2-L2 as a critical determinant of Taf12 function and complex association. This observation aligns with structural data from yeast and human TAFs implicating α2 in defining binding specificity (27, 39). Chimeras involving α1-L1 or α3-L3 showed partial rescue, indicating that other regions within the HFD contribute to stabilizing the heterodimer but may not be essential for initial partner recognition **(Figure 6)**.

### Functional Implications of Taf12 Paralog Divergence

Our findings support a model in which *C. albicans* Taf12 paralogs mediate selective heterodimerization through conserved histone fold domains (HFDs), with the α2-L2 region playing a pivotal role in determining partner specificity. The minimal impact of sequences outside the HFD suggests that core histone fold interactions are sufficient for complex-specific incorporation. However, non-HFD elements such as the conserved Tra1-interacting region (TIR) present exclusively in Taf12L **(Figure S5)** may contribute to complex-specific functions following assembly.

This functional divergence between Taf12 and Taf12L likely reflects evolutionary pressure to partition regulatory roles between TFIID and SAGA, facilitating more refined transcriptional control. Such specialization may enhance *C. albicans’* ability to adapt gene expression programs in response to diverse environmental conditions or host cues an advantage with direct implications for stress tolerance and pathogenicity.

Overall, our findings highlight the modularity of transcriptional coactivator assembly and the capacity of conserved domains to evolve specificity through minor sequence changes. The mechanisms uncovered here may extend to other systems (such as plants) where paralogous subunits confer context-dependent regulatory diversity.

## Materials and methods

### Strains and growth conditions

*C. albicans* strains SN95 were used as parental strain for all strains used here. All strains were cultured in yeast extract peptone-rich medium with either glucose, glucose+doxycycline (50μg/ml) or maltose, as indicated. The genome sequences were obtained from the Candida Genome Database (CGD) for strain construction. Details of strain construction are described in the supplemental text. Strains used in this study are provided in **Table S3**. Plasmids used are listed in **Table S4**. Oligonucleotides used are listed in **Table S5**.

### Immunoblotting

All the indicated strains were cultured in YPM at 30°C for 16-18 hours with shaking at 220 rpm and diluted the culture into fresh YPD with or without doxycycline (50μg/ml) and grown for 8h. Cells were harvested by centrifugation and whole-cell extracts were made by bead beating using chilled glass beads for 8 cycles in lysis buffer as described in (25). Quantified proteins using Bradford assay (Bio-Rad) using BSA as a standard. 100ug was loaded into each well of 10% SDS-PAGE. The blots were probed with anti-TAF12, TAF12L, TAF4, and ADA1 polyclonal antibodies and G6PDH (Sigma). All blots were visualized using the ECL Plus chemiluminescent system.

### Co-Immunoprecipitation

Indicated *C. albicans* strains were cultured in YPM and then diluted to fresh YPD or YPD+ Doxycycline (50ug/ml) with starting O. D600∼0.2 and grown for 8h. Cells were harvested after by centrifugation and whole cell lysate was made as described above.

For TAP Immunoprecipitation: About 5mg equivalent whole cell extract was incubated with 20ul of IgG Sepharose beads for immunoprecipitation. Beads were then washed and incubated overnight with 15U AcTEV protease (invitrogen) and the eluate was directly boiled in an SDS-PAGE sample buffer. 25-50% of immunoprecipitated eluate and 100μg of input samples was separated on 10% SDS-PAGE. The blots were probed with anti-TAF12, TAF12L, TAF4, and ADA1 polyclonal and TAP (invitrogen) antibodies. All blots were visualized using the ECL Plus chemiluminescent system.

For FLAG Immunoprecipitation: About 5mg equivalent whole cell extract was incubated with 15μl of antibody coupled Anti-FLAG M2 Affinity beads (GE) for immunoprecipitation. Beads were then washed and the immunoprecipitated proteins were eluted by once with 100ng and thrice with 200ng of 3xFLAG peptide (Sigma). 25-50% of immunoprecipitated samples and 100μg of input samples was separated on 10% or 4-15% (for HFD experiments) SDS-PAGE. The blots were probed with anti-TAF12, TAF12L, TAF4, and ADA1 polyclonal and FLAG **(Sigma)** antibodies. All blots were visualized using the ECL Plus chemiluminescent system.

### GST Pull-Down assays

For the GST pull-down assay, dilutions of the sonicated whole cell extract from GST only control, GST-Taf4, GST-Ada1, GST-HFDTaf4, GST-HFDAda1, Taf12LFLAG_3_ and Taf12-FLAG_3_ bearing strains were run on an SDS-PAGE gel along with marker dilutions to estimate the amount of fusion protein that was expressed in the lysate. 15μl of Glutathione-sepharose beads were incubated separately with 1nmole, 2nmole and 4nmole equivalent of GST tagged proteins. The beads were washed then incubated with 1nmole of indicated prey proteins.

Bound proteins were eluted by directly boiling the beads and analyzed on either 10% or 10-15% (for HFD only protein) SDS-PAGE gel. The gels were stained with Coomassie Brilliant Blue, followed by multiple rounds of de-staining. The gels were imaged in gel-doc and quantified using ImageJ.

#### Serial dilution assays

*Candida albicans* strains were grown in the YPD liquid media till saturation. The cells were serial diluted for 5, 10-fold dilutions and spotted on indicated plates and incubated at 30°C and imaged.

## Author Contributions

PP and KN conceptualized the study and analyzed the data. PP performed the experiments and PP and KN drafted the manuscript. K.N. acquired funding and supervised the project.

## Author Notes

Conflict of Interest: The author(s) declare no conflict of interest.

## Supplementary information

Table S1

Table S2

Table S3 List of Plasmids.

Table S4 List of Strains.

Table S5 List of Oligonucleotides

Figure S1-S7

Supplementary Figure legends

Supplementary Materials and Methods

