## Supplementary Information for "Histone Fold mediated heterodimerization specifies the selective association of TAF12 paralogs with TFIID and SAGA complexes in *Candida albicans*"

**Supplementary material and methods**

**Cloning and Expression of FLAG tagged *TAF12* variants of *Candida albicans:*** The *CaTAF12L* and *CaTAF12* coding sequences along with 6xHis-2xGly-3xFLAG tag and *ACT1t* sequences were PCR amplified using genomic DNA from strains SKC3 and SKC6 (1) and primers ONC135-ONC943 and ONC137-ONC943 respectively. The PCR products thus obtained were digested sequentially with XbaI-BglII and ligated to NheI-BamHI cut pET28b (+) vector generating plasmids as pPC17 and pPC13 respectively. The plasmids were transformed to BL21 (DE3) bacterial cells and cultured overnight in LB liquid media containing kanamycin. The cells were diluted into fresh LB liquid media with kanamycin. Protein expression was induced with 1mM IPTG and cells were harvested by centrifugation. Cells were resuspended into a buffer containing 40mM HEPES-NaOH pH7.4, 350mM NaCl, 0.1% Tween-20 and 10% Glycerol and sonicated at 10% amplitude for 10s for 3 to 5 number of cycles. The sonicated lysate was cleared by centrifugation at 13000 rpm for 10 minute and soluble supernatant part was collected.

**Cloning and Expression of GST tagged *TAF4* and *ADA1* proteins of *Candida albicans:*** The *CaTAF4* and *CaADA1* coding sequences were PCR amplified using primer pairs ONC151-ONC152 and ONC153-154 respectively and genomic DNA from SC5314 strain as template. The PCR products thus obtained were digested with XhoI and ligated to SmaI-XhoI digested pGEX-5X-3 plasmid vector generating plasmids Ip42 (GST-Taf4) and Ip43 (GST-Ada1). The BL21 bacterial cells expressing these plasmids were cultured overnight in LB liquid media containing ampicillin. The cells were diluted into fresh LB liquid media containing ampicillin. Protein expression was induced with 0.1mM IPTG and cells were harvested by centrifugation. Cells were resuspended into a buffer containing 25mM HEPES-NaOH; pH 7.5, 100mM NaCl, 0.05% NP-40, 5mM DTT and 10% glycerol and sonicated at 10% amplitude for 10s for 3 to 5 number of cycles. The sonicated lysate was cleared by centrifugation at 13000 rpm for 10 minute and soluble supernatant part was collected.

**Cloning and Expression of 6xHis-tagged Histone fold domains (HFDs) of *TAF12* variants of *Candida albicans:*** The gene sequences corresponding to Histone fold domain (HFD) only regions of *TAF12L* (1877-2250 bp) and *TAF12* (1182-1545 bp) were PCR amplified using primer pairs ONC933-ONC934 and ONC935-ONC936, respectively. The PCR products were digested with NcoI-SalI and ligated to similarly cut pET28b (+) vector generating plasmids pPC20

(*HFDTAF12L*-His$_6$) and pPC21 (*HFDTAF12*-His$_6$). The BL21becterial cells expressing these plasmids were cultured overnight in LB liquid media containing kanamycin. The cells were diluted to a starting $OD_{600}$ value of 0.05 into fresh LB liquid media containing kanamycin. Protein expression was induced with 0.1mM IPTG and cells were harvested by centrifugation. Cells were resuspended into a His-binding buffer containing 20mM NaPO4 pH7.4, 500mM NaCl and 20mM Imidazole pH7.4. and sonicated at 10% amplitude for 10s for 3 to 5 number of cycles. The sonicated pellet was solubilized into 8M urea solubilization buffer (His-binding buffer with 8M Urea) at room temperature. The dissolved cell lysate was centrifuged at 13000 rpm for 10 minutes, and the supernatant fraction was collected. This supernatant was then applied to His-Trap 1ml column (GE-Healthcare) pre-equilibrated with 8M urea solubilization buffer. The bound proteins were slowly refolded by applying a linear gradient of 8M urea buffer to no urea His-binding buffer. The refolded proteins were eluted by applying linear gradient of His-elution buffer (20mM NaPO4 pH7.4, 500mM NaCl and 500mM Imidazole pH7.4.). Eluted proteins were concentrated and buffer exchanged to His-binding buffer without imidazole and quantitated using Bio-Rad Bradford reagent.

**Construction of heterozygous *CaADA2* and *CaTAF11* C-terminal TAP tag strains:** *ADA2* and *TAF11* TAP tag strains were generated using the split marker strategy and recyclable *SAT1*-marked tagging cassette (2) from plasmid pPC5. Plasmid pPC5 was generated by ligating a SpeI, end-filled, StuI fragment containing TAP tag along with *ACT1t* terminator from Ip22 to StuI digested plasmid Ip27 (1) containing *SAT1-FLP* marker. We used either *TAF11* specific primers (ONC 948/ONC140 and ONC141/ONC949) or *ADA2* specific primers (ONC 946/ONC140 and ONC141/ONC947) to amplify respective up-split and down-split tagging fragments. The corresponding up and down-split fragments were then transformed to *Candida albicans* strains SN95 (WT), ISC11 ($P_{MAL2}$-*TAF12L*) and ISC12 ($P_{MAL2}$-*TAF12L*) generating heterozygous $Nou^R$ strains SDC24, SDC25 and SDC32 for *TAF11-TAP* respectively and SDC23, SDC26 and SDC31 for *ADA2-TAP* respectively. These $Nou^R$ strains were used to obtain $Nou^S$ segregants as described previously (2) generating strains SDC28, SDC30 and SDC34 for *TAF11-TAP* respectively and SDC27, SDC29 and SDC33 for *ADA2-TAP* respectively.

**Construction of ectopically expressing *TAF12* and *TAF12L* strains:** The $P_{TET}$-*TAF12L-FLAG*$_3$ and $P_{TET}$-*TAF12-FLAG*$_3$ complemented strains were generated as described previously(1). For

$P_{TET}$-*TAF12L-FLAG*$_3$ and $P_{TET}$-*TAF12-FLAG*$_3$ cross complemented strains, plasmids pPC18 bearing *TAF12::3xFLAG-2xGly-6xHis* under the *TET1* promoter and pPC19 bearing *TAF12L::3xFLAG-2xGly-6xHis* under the *TET1* promoter were digested with SacII and Acc65I and transformed to strains ISC11 ($P_{MAL2}$-*TAF12L*) and ISC12 ($P_{MAL2}$-*TAF12L*) respectively. The resultant $Nou^R$ clones were PCR confirmed for correct integration generating strains SDC14 and SDC15 respectively.

For construction of $P_{TET}$-*TAF12-FLAG*$_3$ expressing *taf12*Δ/Δ mutant strain, the *SAT1* marker gene in pPC18 plasmid was replaced by the *ARG4* marker gene as follows. The *ARG4* coding sequence was excised from pSN69 by NotI digestion, end-filled, digested BamHI, and ligated to pPC18 digested with ApaI, end-filled and digested with BglII, and plasmid pPC23 was obtained. Next plasmid pPC23 was digested with SacII and Acc65I and transformed to strains ISC36 (*taf12*Δ/Δ) thereby generating strain SDC18. of $P_{TET}$-*TAF12L-FLAG*$_3$ expressing *taf12*Δ/Δ mutant strain (SDC18) was constructed as described previously (Sinha et al., 2017).

For construction of $P_{MET3}$-*TAF12L-FLAG*$_3$ and $P_{MET3}$-*TAF12-FLAG*$_3$ expressing *taf12*Δ/Δ mutant strain, a PCR fragment encompassing the $P_{MET3}$ sequences was PCR amplified by primers ONC1058-1059 and using plasmid pFA-*CaARG4*-p*MET3* as template. The PCR product thus obtained was digested with NcoI-NdeI and ligated to NcoI-NdeI digested plasmids pPC17 and pPC13 thereby generating plasmids pPC33 and pPC34 respectively. Parallelly, plasmid pCIp10-*CdARG4* was digested with BglII and re-ligated to generate plasmid pPC35. The $P_{MET3}$-*TAF12L-FLAG*$_3$ and $P_{MET3}$-*TAF12-FLAG*$_3$ sequences were excised from plasmids pPC33 and pPC34 as NcoI, end filled and XhoI fragments and ligated to Acc65I, end-filled, XhoI digested pPC35 thereby generating plasmids pPC43 and pPC42. Next these plasmids were digested with StuI and transformed to strains ISC36 (*taf12*Δ/Δ) thereby generating strains SDC35 and SDC36 respectively.

**Construction of ectopically expressing *HFD-TAF12* and *HFD-TAF12L* strains:** Histone fold domain or HFDs of *TAF12L* and *TAF12* along with the 6xHis-2xGly-3xFLAG tag and *ACT1t* sequences were PCR amplified by primers ONC961-ONC943 and ONC942-ONC943 using genomic DNA from strains SKC3 and SKC6 respectively. The resulting PCR products were digested with SalI and BglII and was ligated to similarly cut pNIM1 integrative plasmid (3) thereby generating plasmids pPC12 and pPC11 respectively. These plasmids were then digested with

Acc65I and SacII and the fragments containing *HFD-TAF12L-FLAG3* and *HFD-TAF12-FLAG3* were transformed to ISC11 ($P_{MAL2}$-*TAF12L*) and ISC12 ($P_{MAL2}$-*TAF12*) strains thereby generating strains SDC7, SDC8, SDC2 and SDC5 respectively.

For construction of $P_{TET}$-*HFD-TAF12L-FLAG3* and $P_{TET}$-*HFD-TAF12-FLAG3* expressing *taf12*Δ/Δ mutant strain: *ARG4* gene was popped out from pSN69 plasmid as NotI-end filled-BglII fragment while plasmids pPC12 and pPC11 were digested first with ApaI, end filled and then with BglII removing the *SAT1* marker gene. The two fragments were ligated yielding plasmids pPC14 and pPC15 respectively. These plasmids were digested again with Acc65I and SacII and the fragments containing $P_{TET}$-*HFD-TAF12L-FLAG3* and $P_{TET}$-*HFD-TAF12-FLAG3* were transformed to ISC36 (*taf12l*Δ/Δ) strain, yielding strains SDC10 and SDC11 respectively.

**Construction of *TAF12* depletion strains expressing helix swapped chimeric *TAF12L* constructs:** The helix swapped chimeric *TAF12L* constructs were generated by employing fusion PCR technique (4, 5) and plasmids pPC19 and pPC18 having the native 3XFLAG tagged *TAF12L* and *TAF12* respectively as starting template.

Construction of *TAF12LC1* construct: *TAF12L* α1-L1 swapped chimera was made by fusing three PCR products. Fragment one containing *TAF12L* ORF sequence from internal BmgBI site till the start of α1-Helix of HFD was amplified using the primers ONC1023-ONC1039. A second fragment encompassing *TAF12* ORF sequence of α1-L1 region of HFD was amplified using the primers ONC1040-ONC1041. A third fragment containing *TAF12L* ORF sequence from the start of α2-helix of HFD along with 3XFLAG tag and *Act1* terminator sequences was amplified using primers ONC1042-ONC1022. In a stepwise manner, the three fragments were fused together by fusion PCR and the product thus obtained was then amplified by primers ONC1023-ONC1022, digested with BmgBI and BglII and ligated to BmgBI-BglII cut plasmid pPC31 thereby generating plasmid pPC57.

Construction of *TAF12LC2* construct: *TAF12L* α2-L2 swapped chimera was made by fusing three PCR products. Fragment one containing *TAF12L* ORF sequence from internal BmgBI site till the start of α2-Helix of HFD was amplified using the primers ONC1023-ONC1069. A second fragment encompassing *TAF12* ORF sequence of α2-L2 region of HFD was amplified using the primers ONC1070-ONC1071. A third fragment containing *TAF12L* ORF sequence from the start of α3-helix of HFD along with 3XFLAG tag and *Act1* terminator sequences was amplified using primers ONC1072-ONC1022. In a stepwise manner, the three fragments were fused together by fusion PCR and the product thus obtained was then amplified by primers ONC1023-ONC1022, digested with BmgBI and BglII and ligated to BmgBI-BglII cut plasmid pPC19 thereby generating plasmid pPC58.

Construction of *TAF12LC3* construct: *TAF12L* α3-L3 swapped chimera was made by fusing three PCR products. Fragment one containing *TAF12L* ORF sequence from internal BmgBI site till the start of α3-Helix of HFD was amplified using the primers ONC1023-ONC1073. A second fragment encompassing *TAF12* ORF sequence of α3-L3 region of HFD was amplified using the primers ONC1074-ONC1075. A third fragment containing *TAF12L* ORF sequence from the start of αC-helix of HFD along with 3XFLAG tag and *Act1* terminator sequences was amplified using primers ONC1076-ONC1022. In a stepwise manner, the three fragments were fused together by fusion PCR and the product thus obtained was then amplified by primers ONC1023-ONC1022, digested with BmgBI and BglII and ligated to BmgBI-BglII cut plasmid pPC19 thereby generating plasmid pPC59.

Construction of *TAF12LC4* construct: The *TAF12L* αC swapped chimera was generated by fusing two PCR fragments. One PCR fragment encompassing the *TAF12L* ORF sequence from internal BmgBI site till start of αC of the Histone fold domain (HFD) was amplified using primers ONC1023-ONC1024. A second PCR fragment encompassing the C-terminal α-Helix (αC) of HFD *TAF12* along with 3XFLAG tag and *Act1* terminator sequences was amplified using primers ONC1025-ONC1022. In a stepwise manner, the two fragments were fused together by fusion PCR and the product thus obtained was then amplified by primers ONC1023-ONC1022, digested with BmgBI and BglII and ligated to BmgBI-BglII cut plasmid pNIM1 thereby generating plasmid pPC31.

Construction of *TAF12LC5* construct: The *TAF12L* Histone fold domain (HFD) swapped chimera was generated by fusing two PCR fragments. One PCR fragment encompassing the *TAF12L* ORF sequence from internal BmgBI site till the start of α1-Helix of HFD was amplified using primers ONC1023-ONC1039. A second PCR fragment encompassing the Histone fold domain (HFD) of *TAF12* along with 3XFLAG tag and *Act1* terminator sequences was amplified using primers ONC1040-ONC1022. In a stepwise manner, the two fragments were fused together by fusion PCR and the product thus obtained was then amplified by primers ONC1023-ONC1022, digested with BmgBI and BglII and ligated to BmgBI-BglII cut plasmid pPC31 thereby generating plasmid pPC83.

The plasmids pPC57, pPC58, pPC59, pPC31 and pPC83 were digested with SacII and Acc65I and the fragments containing *TAF12L* chimeric constructs were transformed to strain ISC12 ($P_{MAL2}$-*TAF12*) generating strains SDC41, SDC42, SDC43, SDC21 and SDC55.

**Construction of $P_{TET}$ expressed helix swapped 3XFLAG tagged chimeric *TAF12L* constructs:** To avoid repeated rounds of PCR for construction of *TAF12* helix swapped chimeric constructs, a unique AatII site was introduced in the N-terminal region of *TAF12* by introducing a silent C to G mutation at 1053 bp with respect to ATG as below.

Construction of *TAF12C4* construct: *TAF12* αC-Helix-swapped chimera was made by fusing three PCR products. Fragment one containing *TAF12* ORF sequence from start codon till 1053 bp (till site of introduction of AatII restriction site) was amplified using the primers ONC1017-ONC1018. A second fragment encompassing *TAF12* ORF sequence from site of introduction of AatII restriction site (1054bp) till loop three or L3 of the Histone fold domain (HFD) was amplified using the primers ONC1019-ONC1020. A third fragment encompassing the C-terminal α-Helix (αC) of *TAF12L* along with 3XFLAG tag and *Act1* terminator sequences was amplified using primers ONC1021-ONC1022. In a stepwise manner one, the three fragments were fused together by fusion PCR and the product thus obtained was amplified by primers ONC1017-ONC1022, digested with SalI and BglII and ligated to SalI-BglII cut plasmid pNIM1 thereby generating plasmid pPC32.

Construction of *TAF12C1* construct: *TAF12* α1-L1 swapped chimera was made by fusing three PCR products. Fragment one containing *TAF12* ORF sequence from AatII site till the start of α1-Helix of HFD was amplified using the primers ONC1043-ONC1044. A second fragment encompassing *TAF12L* ORF sequence of α1-L1 region of HFD was amplified using the primers ONC1045-ONC1046. A third fragment containing *TAF12* ORF sequence from the start of α2-helix of HFD along with 3XFLAG tag and *Act1* terminator sequences was amplified using primers ONC1047-ONC1022. In a stepwise manner, the three fragments were fused together by fusion PCR and the product thus obtained was then amplified by primers ONC1043-ONC1022, digested with AatII and BglII and ligated to AatII-BglII cut plasmid pPC32 thereby generating plasmid pPC53.

Construction of *TAF12C2* construct: *TAF12* α2-L2 swapped chimera was made by fusing three PCR products. Fragment one containing *TAF12* ORF sequence from internal AatII site till the start of α2-Helix of HFD was amplified using the primers ONC1043-ONC1077. A second fragment encompassing *TAF12L* ORF sequence of α2-L2 region of HFD was amplified using the primers ONC1078-ONC1079. A third fragment containing *TAF12* ORF sequence from the start of α3-helix of HFD along with 3XFLAG tag and *Act1* terminator sequences was amplified using primers ONC1080-ONC1022. In a stepwise manner, the three fragments were fused together by fusion PCR and the product thus obtained was then amplified by primers ONC1043-ONC1022, digested with AatII and BglII and ligated to AatII-BglII cut plasmid pPC32 thereby generating plasmid pPC67.

Construction of *TAF12C3* construct: *TAF12* α3-L3 swapped chimera was made by fusing three PCR products. Fragment one containing *TAF12* ORF sequence from internal AatII site till the start of α3-Helix of HFD was amplified using the primers ONC1043-ONC1081. A second fragment encompassing *TAF12L* ORF sequence of α3-L3 region of HFD was amplified using the primers ONC1082-ONC1083. A third fragment containing *TAF12* ORF sequence from the start of αC-helix of HFD along with 3XFLAG tag and *Act1* terminator sequences was amplified using primers ONC1084-ONC1022. In a stepwise manner, the three fragments were fused together by fusion PCR and the product thus obtained was then amplified by primers ONC1043-ONC1022, digested with AatII and BglII and ligated to AatII-BglII cut plasmid pPC32 thereby generating plasmid pPC55.

Construction of *TAF12C5* construct: The *TAF12* Histone fold domain (HFD) swapped chimera was generated by fusing two PCR fragments. One PCR fragment encompassing the *TAF12* ORF sequence from internal AatII site till the start of α1-Helix of HFD was amplified using primers ONC1043-ONC1044. A second PCR fragment encompassing the Histone fold domain (HFD) of *TAF12L* along with 3XFLAG tag and *Act1* terminator sequences was amplified using primers ONC1045-ONC1022. In a stepwise manner, the three fragments were fused together by fusion PCR and the product thus obtained was then amplified by primers ONC1043-ONC1022, digested with AatII and BglII and ligated to AatII-BglII cut plasmid pPC32 thereby generating plasmid pPC56.

The plasmids pPC32, pPC53, pPC67, pPC55 and pPC56 were digested with SacII and Acc65I and the fragments containing *TAF12* chimeric constructs transformed to strain ISC12 ($P_{MAL2}$-*TAF12*) generating strains SDC22, SDC51, SDC54, SDC52 and SDC44.

**Table S1: List of amino-acids markedly different between two CaTaf12 variants**

| CaTaf12L | CaTaf12 | yTaf12 | hTaf12 |
|---|---|---|---|
| N638 | R407 | S416 | T59 |
| T648 | N417 | T426 | E69 |
| G650 | S419 | G428 | - |
| E653 | Q422 | E431 | - |
| T658 | I427 | T436 | E79 |
| G662 | N431 | G440 | G78 |
| N663 | D432 | D441 | D79 |
| L668 | F437 | L446 | L84 |
| H677 | R446 | T455 | E93 |
| S678 | N447 | N456 | S94 |
| T680 | V449 | T458 | V96 |
| A683 | S452 | S461 | A99 |
| C684 | G453 | C462 | C100 |
| S694 | R463 | N472 | T110 |
| H703 | N472 | H481 | H119 |
| N709 | G478 | N487 | N125 |
| M717 | T486 | A495 | S133 |
| T723 | A492 | T501 | Y130 |
| S731 | E500 | N509 | A147 |
| Q734 | E503 | Q512 | Q150 |
| E740 | S509 | T518 | R156 |
| V745 | N512 | A523 | S159 |
| N746 | K513 | A524 | K160 |

**Table.S2. Summary of Class specific residues of CaTaf12 variants and respective amino-acids of CaTaf4 and CaAda1 involved in protein-protein interactions.**

| CaTaf12L | CaAda1 | CaTaf4 | CaTaf12 | CaAda1 | CaTaf4 |
|---|---|---|---|---|---|
| N638 | I233<br>D235 | H136 | T407 | I233 | H136 |
| T648 | - | - | N417 | - | - |
| G650 | - | - | S419 | V282<br>I280 | - |
| E653 | - | - | Q422 | A279<br>I280 | K181<br>T182<br>I183<br>I184 |
| T658 | - | - | I427 | - | - |
| G662 | - | - | N431 | - | - |
| N663 | - | - | D432 | - | - |
| L668 | - | L175 | F437 | I275 | L178 |
| H677 | M242 | F143 | R446 | - | - |
| S678 | - | - | N447 | - | - |
| T680 | M264<br>M247<br>I246<br>M243 | V147<br>N151 | V449 | M264<br>M247<br>M243 | V147<br>N151 |
| A683 | - | - | S452 | - | - |
| C684 | H250<br>M247 | V147<br>A148<br>N151 | G453 | L252 | N151 |
| S694 | L252<br>T253<br>G254<br>G255 | I153 | R463 | L256<br>G255<br>G254<br>T253<br>L252 | F157<br>L158<br>M159 |
| H703 | - | - | N472 | - | - |
| N709 | - | - | G478 | - | - |

#### Table S3: List of Plasmids

| PLASMID | RELEVANT DESCRIPTION | SOURCE |
|---|---|---|
| pNIM1 | *C.a. SAT1 CaADH1* p*Tet-CaGFP* | (6) |
| pLITMUS28 | Cloning vector, $Amp^R$ | NEB |
| pET28b (+) | T7 lac promoter with N- and C-terminal 6xHis tag | Novagen |
| pGEX-5X-3 | $P_{tac}$promoter with N-terminal GST tag | GE Healthcare |
| pCIp10-*C.d. ARG4* | pCIp10 with *C.d. ARG4* in place of *C.a.URA3* | (1) |
| pFA-*CaARG4-pMET3* | | |
| pSN69 | *C.d. ARG4* | (6) |
| Ip22 | TAP tag, *C.d. HIS1* | (1) |
| Ip27 | *SAT1FLP* cassette in pLITMUS28 | (1) |
| Ip42 | *TAF4* in pGEX-5X-3 vector | (1) |
| Ip43 | *ADA1* in pGEX-5X-3 vector | (1) |
| Ip50 | 6xHis-2xGly-3xFLAG tag, *ACT1*t, with *SAT1 FLP* | (1) |
| pPC5 | TAP tag, *ACT1*t, *SAT1 FLP* in pLITMUS28 vector | This work |
| pPC11 | *TAF12* (1182-1545), 6xHis-2xGly-3xFLAG in pNIM1 vector | This work |
| pPC12 | *TAF12L* (1877-2250bp), 6xHis-2xGly-3xFLAG in pNIM1 vector | This work |
| pPC13 | *TAF12*, 6xHis-2xGly-3xFLAG in pET28b (+) vector | This work |
| pPC14 | *TAF12L* (1877-2250bp), 6xHis-2xGly-3xFLAG in pNIM1 vector bearing *ARG4* instead of *SAT1* | This work |
| pPC15 | TAF12 (1182-1545), 6xHis-2xGly-3xFLAG in pNIM1 vector bearing *ARG4* instead of *SAT1* | This work |
| pPC16 | pNIM1 vector bearing *ARG4* instead of *SAT1* | (1) |
| pPC17 | *TAF12L*, 6xHis-2xGly-3xFLAG in pET28b (+) vector | This work |
| pPC18 | *TAF12*, 6xHis-2xGly-3xFLAG in pNIM1 vector | (1) |
| pPC19 | *TAF12L*, 6xHis-2xGly-3xFLAG in pNIM1 vector | (1) |
| pPC20 | HFD of *TAF12L* in pET28b (+) vector | This work |
| pPC21 | HFD of *TAF12* in pET28b (+) vector | This work |
| pPC22 | *TAF12L*, 6xHis-2xGly-3xFLAG in pNIM1 vector bearing *ARG4* instead of *SAT1* | (1) |
| pPC23 | *TAF12*, 6xHis-2xGly-3xFLAG in pNIM1 vector bearing *ARG4* instead of *SAT1* | This work |
| pPC28 | HFD of *ADA1* in pGEX-5X-3 vector | This work |
| pPC29 | HFD of *TAF4* in pGEX-5X-3 vector | This work |
| pPC31 | *TAF12L*C4, 6xHis-2xGly-3xFLAG in pNIM1 vector | This work |
| pPC32 | *TAF12*C4, 6xHis-2xGly-3xFLAG in pNIM1 vector | This work |
| pPC33 | pPC17 with $P_{MET3}$ upstream of *TAF12L* coding sequence | This work |
| pPC34 | pPC13 with $P_{MET3}$ upstream of *TAF12* coding sequence | This work |

| | | |
|---|---|---|
| pPC35 | pCIp10-*C.d. ARG4* BglII site inactivated | This work |
| pPC42 | $P_{MET3}$-*TAF12* in pPC35 | This work |
| pPC43 | $P_{MET3}$-*TAF12L* in pPC35 | This work |
| pPC53 | *TAF12*C1 fragment, 6xHis-2xGly-3xFLAG in pPC32 | This work |
| pPC54 | *TAF12*C2 fragment, 6xHis-2xGly-3xFLAG in pPC32 | This work |
| pPC55 | *TAF12*C3 fragment, 6xHis-2xGly-3xFLAG in pPC32 | This work |
| pPC56 | *TAF12*C5 fragment, 6xHis-2xGly-3xFLAG in pPC32 | This work |
| pPC57 | *TAF12L*C1 fragment, 6xHis-2xGly-3xFLAG in pPC31 | This work |
| pPC58 | *TAF12L*C2 fragment, 6xHis-2xGly-3xFLAG in pPC31 | This work |
| pPC59 | *TAF12L*C3 fragment, 6xHis-2xGly-3xFLAG in pPC31 | This work |
| pPC62 | 6xHis-2xGly-3xFLAG tag in pNIM1 vector | This work |
| pPC66 | *TAF12LC5* fragment, 6xHis-2xGly-3xFLAG in pPC31 | This work |
| pPC67 | *TAF12*C2 fragment, 6xHis-2xGly-3xFLAG in pPC32 | This work |

#### Table S4: List of Strains

| STRAINS | RELEVANT GENOTYPE | SOURCE |
|---|---|---|
| SN95 | *arg4Δ/arg4Δ his1Δ/his1Δ URA3/ura3Δ::*$imm^{434}$ *IRO1/iro1Δ::*$imm^{434}$ | (6) |
| ISC11 | SN95 *HAH1-*$P_{MAL2}$*-TAF12L/HIS1-*$P_{MAL2}$*-TAF12L* | (1) |
| ISC12 | SN95 *HAH1-*$P_{MAL2}$*-TAF12/HIS1-*$P_{MAL2}$*-TAF12* | (1) |
| ISC36 | SN95 *taf12l Δ::FRT/ taf12lΔ::SAT1 FLP* | (1) |
| SKC3 | SN87 *TAF12L::His6-Gly2-FLAG3-FRT/TAF12 L::His6-Gly2-FLAG3-SAT1-FLP* | (1) |
| SKC6 | SN87 *TAF12:: His6-Gly2-FLAG3-FRT/TAF12 ::His6-Gly2-FLAG3-SAT1-FLP* | (1) |
| SDC2 | ISC11 *ADH1/adh1*<pPC11> | This work |
| SDC3 | ISC11 *ADH1/adh1*<pNIM1> | (1) |
| SDC5 | ISC12 *ADH1/adh1*<pPC11> | This work |
| SDC6 | ISC12 *ADH1/adh1*<pNIM1> | (1) |
| SDC7 | ISC11 *ADH1/adh1*<pPC12> | This work |
| SDC8 | ISC12 *ADH1/adh1*<pPC12> | This work |
| SDC9 | SN95 *taf12lΔ::FRT/ taf12lΔ::FRT* | This work |
| SDC10 | ISC36 *ADH1/adh1*<pPC14> | This work |
| SDC11 | ISC36 *ADH1/adh1*<pPC15> | This work |
| SDC12 | ISC36 *ADH1/adh1*<pPC16> | (1) |
| SDC13 | ISC11 *ADH1/adh1*<pPC19> | (1) |
| SDC14 | ISC11 *ADH1/adh1*<pPC18> | This work |
| SDC15 | ISC12 *ADH1/adh1*<pPC19> | This work |
| SDC16 | ISC12 *ADH1/adh1*<pPC18> | (1) |
| SDC17 | ISC36 *ADH1/adh1*<pPC22> | (1) |

| | | |
|---|---|---|
| SDC18 | ISC36 *ADH1/adh1*<pPC23> | This work |
| SDC21 | ISC12 *ADH1/adh1*<pPC31> | This work |
| SDC22 | ISC12 *ADH1/adh1*<pPC32> | This work |
| SDC23 | SN95 *ADA2::TAP-SAT1 FLP/ADA2* | This work |
| SDC24 | SN95 *TAF11::TAP-SAT1 FLP/TAF11* | This work |
| SDC25 | ISC11 *TAF11::TAP-SAT1 FLP/TAF11* | This work |
| SDC26 | ISC11 *ADA2::TAP-SAT1 FLP/ADA2* | This work |
| SDC27 | SN95 *ADA2::TAP-FRT/ADA2* | This work |
| SDC28 | SN95 *TAF11::TAP-FRT/TAF11* | This work |
| SDC29 | ISC11 *ADA2::TAP-FRT/ADA2* | This work |
| SDC30 | ISC11 *TAF11::TAP-FRT/TAF11* | This work |
| SDC31 | ISC12 *ADA2::TAP-SAT1 FLP/ADA2* | This work |
| SDC32 | ISC12 *TAF11::TAP-SAT1 FLP/TAF11* | This work |
| SDC33 | ISC12 *ADA2::TAP-FRT/ADA2* | This work |
| SDC34 | ISC12 *TAF11::TAP-FRT/TAF11* | This work |
| SDC35 | ISC36 *RPS10/rps10*<pPC43> | This work |
| SDC36 | ISC36 *RPS10/rps10*<pPC42> | This work |
| SDC37 | SDC33 *ADH1/adh1*<pPC57> | This work |
| SDC38 | SDC33 *ADH1/adh1*<pPC58> | This work |
| SDC39 | SDC33 *ADH1/adh1*<pPC59> | This work |
| SDC40 | SDC33 *ADH1/adh1*<pPC56> | This work |
| SDC41 | SDC34 *ADH1/adh1*<pPC57> | This work |
| SDC42 | SDC34 *ADH1/adh1*<pPC58> | This work |
| SDC43 | SDC34 *ADH1/adh1*<pPC59> | This work |
| SDC44 | SDC34 *ADH1/adh1*<pPC56> | This work |
| SDC49 | SDC33 *ADH1/adh1*<pPC53> | This work |
| SDC50 | SDC33 *ADH1/adh1*<pPC55> | This work |
| SDC51 | SDC34 *ADH1/adh1*<pPC53> | This work |
| SDC52 | SDC34 *ADH1/adh1*<pPC55> | This work |
| SDC53 | SDC33 *ADH1/adh1*<pPC67> | This work |
| SDC54 | SDC34 *ADH1/adh1*<pPC67> | This work |
| SDC55 | SDC34 *ADH1/adh1*<pPC66> | This work |

#### Table S5: List of Oligonucleotides

| Primer | SEQUENCE (5’ to 3') | NOTES |
|---|---|---|
| ONC103 | AATCTATTACTCAATCGAG | 5’-primer for diagnostic PCR of pNIM1 integration; binds to $P_{TET}$ region. |
| ONC104 | TAAAAATATCGCACTCAC | 3’-primer for *ADH1;* +1374 to +1356 wrt ATG |
| ONC136 | GCCTCGAGAGCATTATCATCATTCAC | Reverse primer for *ORF 19.470* with XhoI site (Position +2256 to +2239 wrt ATG) |
| ONC137 | GGTCTAGAATGGATTCATCAGCAGCT | Forward primer for *ORF 19.6820* with XbaI site for in frame insertion into pET 28b+ and 2 bp overhang for in frame insertion ino pGEX-5x-3 (Position +1 to +18 wrt ATG) |

| | | |
|---|---|---|
| ONC140 | TCTTGGTGAGAACAGCGACCGAAA | Reverse primer for upstream split marker of SAT1 flipper |
| ONC141 | GGAGCGATAAGCGTGCTTCTGCCG | Forward primer for downstream split marker of SAT1 flipper |
| ONC151 | gcgctagcATGACAAGTACACCTCAA | Forward primer for *CaTAF4* ORF with NheI site for in frame insertion into pET 28b+ and 2 bp overhang for in frame insertion ino pGEX-5X-3. +1 to +18 wrt ATG |
| ONC152 | gctctcgagATCTTTCAATTTTGCATA | Reverse primer for *CaTAF4* ORF with XhoI site +1089 to +1072 wrt ATG |
| ONC153 | gcgctagcATGACATCTCAAATCGCT | Forward primer for *CaADA1* ORF with NheI site for in frame insertion into pET 28b+ and 2 bp overhang for in frame insertion ino pGEX-5X-3. +1 to +18 wrt ATG |
| ONC154 | gctctcgagCATAGTCGACACTAAATC | Reverse primer for *CaADA1* ORF with XhoI site +1470 to +1453 wrt ATG |
| ONC845 | CGCTCGAGGGTGCTTTTTTCCAA | Reverse primer fot*TAF4* HF cloning with 5'-GC 2bp overhang and XhoI site |
| ONC847 | CCCTCGAGCGATTCGACTAAATGA | Reverse primer fot*ADA1* HF cloning with 5'-GG 2bp overhang and XhoI site |
| ONC933 | CCCCATGGTTATGATTGATTCTAGTACCAGT GGA | FORWARD primer for HF-*TAF12L* cloning with ATG codon and NcoI site and 2bp overhang at 5'end |
| ONC934 | GCCTCGAGTTAGTCGACAGCATTATCATCA TTCACA | Reverse primer for HF-*TAF12L* cloning with TAA stop codon between SalI and XhoI site and 2bp overhang at 3'end |
| ONC935 | GTCCATGGTTATGAACATACCCGACAA TGATG | Forward primer for HF-*TAF12* cloning with ATG codon and NcoI site and 2 bp overhang at 5'end |
| ONC936 | GTCTCGAGCTAGTCGACTTGATCTTTATTTA ATTTGGA | Reverse primer for HF-*TAF12* cloning with TAA stop codon between SalI and XhoI site and 2bp overhang at 3'end |
| ONC937 | GAGCTAGCATGGTGATCAGGTCCAGCAAAC T | Forward primer for HF-*TAF4* cloning with ATG codon and NheI site and 2bp overhang at 5'end |
| ONC938 | GAGCTAGCATGATAAATACACCAATAGCCA CTGAA | Forward primer for HF-*ADA1* cloning with ATG codon and NheI site and 2bp overhang at 5'end |
| ONC939 | GCCGTCGACATGAATAATGGATCTCAGAAT C | Forward Primer for full length *TAF12L* cloning for FL complementation with SalI site, ATG codon and 3bp 5' overhang |
| ONC940 | GGAGTCGACATGGATTCATCAGCAGCTTC | Forward Primer for full length *TAF12* cloning for FL complementation with SalI site, ATG codon and 3bp 5' overhang |
| ONC942 | GGAGTCGACATGCCAAGTAACATACCCGAC AAT | Forward Primer for *TAF12* HFD cloning for HF complementation with SalI site, ATG codon and 3bp 5' overhang |
| ONC943 | GGAAGATCTCCCCGAAGATCTTATGATGGA | Reverse Primer for *TAF12L*/*12b* HF complementation in the FLAG-tag Act1 terminator region with BglII site and 3bp overhangs |

| | | |
|---|---|---|
| ONC946 | GTCAATAAAGCTTCTAAAATCTATGAATTCTTTGTGCATATGGGATGGTGTTCTCAGGGGGGATCCATGGAAAAGAGAAG | Forward Primer for *ADA2*-TAP tagging |
| ONC947 | GATCCACTAG**CCT**TGACTAGATGTAGTATATTCTTTTCATTTTTTTTGTTTTGCATAGATTTATATAATTGGATTTAAA | Reverse Primer for *ADA2*-TAP tagging |
| ONC948 | AACAGTGTATTTTCTGGAAGTCGGAAAAGAAAAATGGGAGACGATGGCCCTTTCTATGTTGGATCCATGGAAAAGAGAAG | Forward Primer for *TAF11*-TAP tagging |
| ONC949 | AACAAACCAGAATACTGCCTAAGAAATACAAAACAAAACTAAGCATTAGCAAAGCCATAACTAGTCAAGGCTAGTGGATC | Reverse Primer for *TAF11*-TAP tagging |
| ONC961 | GGAGTCGACATGGGTGTCATTGATTCTAGTACCA | Forward Primer for *TAF12L* HFD cloning for HF complementation with SalI site, ATG codon and 3bp 5' overhang |
| ONC973 | CATGGTACCGTCGACATGTCAAAGGATTC | Forward primer with KpnI (Acc65I) site and 3bp overhangs to clone *ADH1p* to CIp10 |
| ONC974 | TCCCTCGAGCATAATTGTTTTTGTATTTG | Reverse primer with XhoI site and 3bp overhangs to clone *ADH1p* to CIp10 |
| ONC1017 | ACACATACAAATATAAATAGTCGACATGGATTCATCAGCAGCTTC | Forward primer amplifying *TAF12* N-terminal region till 1052 bp (before and excluding the AatII site mutation) and overlapping pNIM1 backbone. |
| ONC1018 | AGCTGACGTCGATGTTGTCGCAGTAAGTGG | Reverse primer amplifying *TAF12* N-terminal part at 1052 bp (before and excluding the AatII site mutation) and overlapping the *TAF12* N-terminal region at 1053 bp (after and including AatII site mut). |
| ONC1019 | CGACAACATCGACGTCAGCTGTGGCACC | Forward primer amplifying the *TAF12* N-terminal region from 1053 bp (after and including AatII site mut) and overlapping *TAF12* N-terminal region till 1052 bp (before and excluding the AatII site mutation) |
| ONC1020 | GTATTTTTCTTGTTGCTCTTATTTCATCAGTAGAATATCC | Reverse primer amplifying the *TAF12* N-terminal part at 1052 bp (after and including AatII site mut) and overlappingTAF12L αC region |
| ONC1021 | AATAAGAGCAACAAGAAAAATACAACCTAGTAATAGTTATAG | Forward primer amplifying the *TAF12L* αC region and overlapping the *TAF12*-Nterminal part at 1052 bp (after and including AatII site mut) |
| ONC1022 | TTCCAGATTTCCAGAATTTAGATCTATGATGGAATGAATGGGATG | Reverse primer amplifying the *TAF12L* αC region and overlapping pNIM1 backbone Common Reverse primer for each construct-amplifying *TAF12L* or *TAF12* αC region and overlapping pNIM1/pPC18/pPC19 backbone. |
| ONC1023 | GAATTTGGCTGGCATTACACGTCAGTCTGTCCCATCATTGC | Forward primer amplifying *TAF12L* N-terminal part after the BmgBI site and overlapping pPC19 (in *TAF12L* ORF region before BmgBI site). |
| ONC1024 | ATTTTCTTGCGTTTCGAATCTCATCCATTG | Reverse primer amplifying *TAF12L* N-terminal part (after the BmgBI site) and overlapping TAF12 αC region |

| | | |
|---|---|---|
| ONC1025 | GATTCGAAACGCAAGAAAATGGCAACCTG | Forward primer amplifying *TAF12* αC region and overlapping *TAF12L* N-terminal part (after the BmgBI site) |
| ONC1039 | TCTTAGTCAACACTCGTCCACTGGTACTAG AATC | Reverse overlapping primer to amplify the N-terminal region of *TAF12L* before 12aC1 RP1 12aC5 RP1 |
| ONC1040 | AGTACCAGTGGACGAGTGTTGACTAAGAGA AAGTTAGTGG | Forward primer with a 3' annealing region complementary to the α1-Helix of TAF12 while a 5' region complementary to TAF12L sequence just before the start of α1 helix of HFD |
| ONC1041 | CCTCCACATTCCCATCAATAGGAATTTTGGC ATCTC | Reverse primer with a 3' annealing region complementary to the loop L1 of HFD of TAF12 while a 5' region complementary toTAF12L sequence just after the loop L1 of HFD |
| ONC1042 | CAAAATTCCTATTGATGGGAATGTGGAGGA AT | Forward primer with a 3' annealing region complementary to TAF12L sequence just after the loop L1 helix of HFD while a 5' region complementary to the α1-Helix of TAF12 |
| ONC1043 | GACAACATCGACGTCAGCTGTGGCACCA | Forward primer with a 3' annealing region complementary to the α1-Helix of TAF12L while a 5' region complementary to TAF12 sequence just before the start of α1 helix of HFD |
| ONC1044 | GTTTATTTAAAACTCTGCCATCATTGTCGGG | Reverse primer with a 3' annealing region complementary to TAF12 sequence just before the start of α1 helix of HFD while a 5' region complementary to the α1-Helix of TAF12L |
| ONC1045 | CCCGACAATGATGGCAGAGTTTTAAATAAA CGAAAATTAGGTG | Forward primer with a 3' annealing region complementary to the α1-Helix of TAF12L while a 5' region complementary to TAF12 sequence just before the start of α1 helix of HFD |
| ONC1046 | ATCTTCAACATCGTTATCAATACTGGTCTTA CCATCCC | Reverse primer with a 3' annealing region complementary to the α1-Helix of TAF12L while a 5' region complementary to TAF12 sequence just after the loop L1 helix of HFD. |
| ONC1047 | TAAGACCAGTATTGATAACGATGTTGAAGA TATTTTC | Forward primer with a 3' annealing region complementary to TAF12L sequence just after the loop L1 helix of HFD while a 5' region complementary to the α1-Helix of TAF12 |
| ONC1069 | CAACATCGTTATCAATACTGGTCTTACCATC CC | Forward primer a 3' annealing region complementary to TAF12L sequence just before the start of α2 helix of HFD while a 5' region complementary to the α2-Helix of TAF12 |
| ONC1070 | TAAGACCAGTATTGATAACGATGTTGAAGA TATTTTC | Forward primer with a 3' annealing region complementary to the α2-Helix of TAF12 while a 5' region complementary toTAF12L sequence just before the start of α2 helix of HFD |

| | | |
|---|---|---|
| ONC1071 | CATCTCTTGCCTCAATTCTATCCAATTTTCT ATGTTTG | Reverse primer with a 3' annealing region complementary to the L2 loop of TAF12 HFD while a 5' region complementary toTAF12L sequence just after the loop L2 |
| ONC1072 | ATTGGATAGAATTGAGGCAAGAGATGTTCA AC | Forward primer with a 3' annealing region complementary to TAF12L sequence just after the loop L2 of HFD while a 5' region complementary to the loop L2 of TAF12 HFD |
| ONC1073 | CATCTCTTACGTCTATACTATCCACCTTTCT ATGTTTTG | Reverse primer with a 3' annealing region complementary to TAF12L sequence just before the start of α3 helix of HFD while a 5' region complementary to the α3-Helix of TAF12 |
| ONC1074 | AAGGTGGATAGTATAGACGTAAGAGATGTG CAATTAAATTTAG | Forward primer with a 3' annealing region complementary to the α3-Helix of TAF12 while a 5' region complementary to TAF12L sequence just before the start of α3 helix of HFD |
| ONC1075 | TTGTATTTTTCTTGTTGCTCTTATTTCATCAG TAGAATATC | Reverse primer with a 3' annealing region complementary to the L3 loop of TAF12 HFD while a 5' region complementary to TAF12L sequence just after the loop L3 |
| ONC1076 | TGAAATAAGAGCAACAAGAAAAATACAAC CTAGTAATAGTTATAG | Forward primer with a 3' annealing region complementary to TAF12L sequence just after the loop L3 of HFD while a 5' region complementary to the loop L3 of TAF12 HFD |
| ONC1077 | CCACATTCCCATCAATAGGAATTTTGGCAT CTCCTTG | Reverse primer with a 3' annealing region complementary to TAF12 sequence just before the start of α2 helix of HFD while a 5' region complementary to the α2-Helix of TAF12L |
| ONC1078 | CAAAATTCCTATTGATGGGAATGTGGAGGA ATT | Forward primer with a 3' annealing region complementary to the α2-Helix of TAF12L while 5' region complementary to TAF12 sequence just before the start of α2 helix of HFD |
| ONC1079 | CATCTCTTACGTCTATACTATCCACCTTTCT ATGTTTTGCTAAC | Reverse primer with a 3' annealing region complementary to the Loop L2 of TAF12L HFD while a 5' region complementary toTAF12 sequence just after the loop L2 of HFD |
| ONC1080 | GGTGGATAGTATAGACGTAAGAGATGTGCA ATTAAATTTAG | Forward primer with a 3' annealing region complementary to TAF12 sequence just after the loop L2 of HFD while a 5' region complementary to the loop L2 of TAF12L |
| ONC1081 | CATCTCTTGCCTCAATTCTATCCAATTTTCT ATGTTTGGC | Reverse primer with a 3' annealing region complementary to TAF12 sequence just before the start of α3 helix of HFD while a 5' region complementary to the α3-Helix of TAF12L. |
| ONC1082 | AAATTGGATAGAATTGAGGCAAGAGATGTT CAAC | Forward primer with a 3' annealing region complementary to the α3-Helix of TAF12L while a 5' region complementary to TAF12 |

| | | |
|---|---|---|
| | | sequence just before the start of α3 helix of HFD |
| ONC1083 | TTGCCATTTTCTTGCGTTTCGAATCTCATCC ATTG | Reverse primer with a 3' annealing region complementary to the Loop L3 of TAF12L HFD while a 5' region complementary to TAF12 sequence just after the loop L3 of HFD |
| ONC1084 | TGAGATTCGAAACGCAAGAAAATGGCAAC CTG | Forward primer with a 3' annealing region complementary to TAF12 sequence just after the loop L3 of HFD while a 5' region complementary to the loop L3 of HFD of TAF12L |
| ONC1104 | TCTCGAGCATCATCACCATCAT | Forward primer FLAG to clone to pNIM1 vector |
| ONC1105 | GGCACGATATCTTATTTATCATCATCATC | Reverse primer FLAG to clone to pNIM1 vector |

#### Supplementary Figure legends

**Figure S1. Non-redundant association of CaTaf12 variants. (A-C)** Western blot confirmation of TAP tagged *CaADA2* and *CaTAF11* strains. All the strains were grown for 7h in YPD liquid media at 30°C and 220rpm and whole cell extracts were made, run on 10% SDS PAGE gel and blotted to nitrocellulose membranes. The membranes were probed with anti-TAP (1:3000) and anti-G6PDH (1:3000). **(A)** Lane 1: WT untagged (SN95), lanes 2-3: two serial dilutions of Ada2-TAP in WT (SDC27) and lanes 4-5: two serial dilutions of Taf11-TAP in WT (SDC28) **(B)** Lane 1: $P_{MAL2}$-*TAF12L* (ISC11), lanes 2-3: two serial dilutions of Ada2-TAP in $P_{MAL2}$-*TAF12L* (SDC29) and lanes 4-5: two serial dilutions of Taf11-TAP in $P_{MAL2}$-*TAF12L* (SDC30). **(C)** Lane 1: WT untagged (SN95), lane 2: $P_{MAL2}$-*TAF12* (ISC12), lanes 3-4: two replicates of Ada2-TAP in $P_{MAL2}$-*TAF12* (SDC33) and lanes 5-6: two replicates of Taf11-TAP in $P_{MAL2}$-*TAF12* (SDC34). **(D)** Experimental strategy used for genetic analysis of functional redundancy of Ca*TAF12* variants. (i-ii) $P_{MAL2}$-*TAF12* (ISC12) strain expressing (i) $P_{TET}$-*TAF12* or (ii) $P_{TET}$-*TAF12L*. (iii-iv) $P_{MAL2}$-*TAF12L* (ISC11) strain expressing (iii) $P_{TET}$-*TAF12L* or (iv) $P_{TET}$-*TAF12*. **(E-F)** Western blot confirmation of full length *TAF12L* and *TAF12* complemented and cross-complemented strains from (D)**.** All the strains were grown in YPM liquid media for 14-16h. The cultures were diluted to fresh YPD+Doxycycline (50µg/ml) media and grown for 7h. Whole cell extracts were made, run on 10% SDS PAGE gel and blotted to nitrocellulose membranes. The membranes were probed with anti-FLAG (1:1000) and anti-G6PDH (1:3000) antibodies. **(E)** Lane 1: *TAF12*-$FLAG_3$ in WT (SKC6), lane 2: WT untagged (SN95), lane 3: $P_{MAL2}$-*TAF12* (ISC12), lane 4-6: three serial dilutions for $P_{TET}$ expressed *TAF12L*-$FLAG_3$ in ISC12 (SDC15) and lanes 7-9: three serial dilutions for $P_{TET}$ expressed *TAF12*-$FLAG_3$ in ISC12 (SDC16). **(F)** Lane 1: *TAF12L*-$FLAG_3$ in WT (SKC3), lane 2: WT untagged (SN95), lane 3: $P_{MAL2}$-*TAF12L* (ISC11), lane 4-6: three serial dilutions for $P_{TET}$ expressed *TAF12L*-$FLAG_3$ in ISC11 (SDC13) and lanes 7-9: three serial dilutions for $P_{TET}$ expressed *TAF12*-$FLAG_3$ in ISC11 (SDC14). Asterisk (*) represents Non-specific bands.

**Figure S2. Ectopically *$P_{TET}$* expressed CaTaf12 variants cannot complement the absence of the other variant.** **(A)** Western blot confirmation of *$P_{TET}$* expressed full length *TAF12L* and *TAF12* in *taf12lΔ/Δ* strains. All the strains were grown in YPD+doxycycline liquid media at 30°C and 220rpm and whole cell extracts were made, run on 10% SDS PAGE gel and blotted to nitrocellulose membranes. The membranes were probed with anti-FLAG (1:1000), anti-TAF12L (1:1000) and anti-G6PDH (1:3000) antibodies. Lane 1: *TAF12L*-$FLAG_3$ in WT (SKC3), lane 2: *taf12lΔ/Δ* (ISC36), lane 3: $P_{TET}$ expressed *TAF12L*-$FLAG_3$ in *taf12l*Δ/Δ (SDC17) and lane 4: $P_{TET}$ expressed *TAF12*-$FLAG_3$ in *taf12l*Δ/Δ (SDC18). Asterisk (*) represents non-specific bands. **(B)** Effect of ectopic expression of *TAF12* and *TAF12L* from *TET* promoter on growth of *taf12l*Δ/Δ mutant. WT strain (SN95), *taf12l*Δ/Δ (ISC36), $P_{TET}$-*TAF12L*-$FLAG_3$ in *taf12l*Δ/Δ (SDC17), $P_{TET}$-*TAF12*-$FLAG_3$ in *taf12l*Δ/Δ (SDC18) and vector only control (SDC12) was grown in YPD liquid media till saturation and then serial dilutions were spotted on YPD, YPD+Doxycycline (50μg/ml), YPD+menadione (90μM) and YPD+Doxycycline+menadione plates. **(C)** Western blot confirmation of *$P_{MET3}$* expressed full length *TAF12L* and *TAF12* in *taf12lΔ/Δ* strains. All the strains were grown in SC liquid media with and without Methionine and Cysteine at 30°C and 220rpm and whole cell extracts were made, run on 10% SDS PAGE gel and blotted to nitrocellulose membranes. The membranes were probed with anti-FLAG (1:1000) and anti-G6PDH (1:3000) antibodies. Lane 1 and 7: WT (SN95), lane 2 and 8: *taf12lΔ/Δ* (ISC36), lane 3-4 and 9-10: two biological replicates of $P_{MET3}$ expressed *TAF12L*-$FLAG_3$ in *taf12l*Δ/Δ (SDC35) and lane 5-6 and 11-12: two biological replicates of $P_{MET3}$ expressed *TAF12*-$FLAG_3$ in *taf12l*Δ/Δ (SDC36). Asterisk (*) represents Non-specific bands. **(D)** Effect of ectopic expression of *TAF12* and *TAF12L* from *MET3* promoter on growth of *taf12l*Δ/Δ mutant. WT strain (SN95), *taf12l*Δ/Δ (ISC36), $P_{MET3}$-*TAF12L*-$FLAG_3$ in *taf12l*Δ/Δ (SDC35), $P_{TET}$-*TAF12*-$FLAG_3$ in *taf12l*Δ/Δ (SDC36) were grown in SC+Methionine+cysteine liquid media till saturation and then serial dilutions were spotted on SC+Met+Cys and SC-Met-Cys plates. **(E)** Co-immunoprecipitation analysis of *$P_{TET}$* expressed

Taf12L and Taf12 in the absence of Taf12. 5mg equivalent of Whole cell protein extracts prepared from 7h YPD+Doxycycline grown strains *TAF12*-$FLAG_3$ (SKC6), *TAF12L*-$FLAG_3$ (SKC3), $P_{MAL2}$-*TAF12* (ISC12), $P_{TET}$-*TAF12L*-$FLAG_3$ in ISC12 (SDC15) and $P_{TET}$-*TAF12*-$FLAG_3$ in ISC12 (SDC16) were immunoprecipitated with FLAG-M2 affinity gel and eluted using 3XFLAG peptide. 100μg equivalent protein extract from each strain was used as input while 25% of the total IP elute was used. Membrane was probed with anti-FLAG (1:1000), anti-Ada1 (1:1000) and anti-Taf4 (1:3000). Lane 1-4 Input: $P_{MAL2}$-*TAF12* (ISC12), *TAF12*-$FLAG_3$ (SKC6), $P_{TET}$-*TAF12L*-$FLAG_3$ in ISC12 (SDC15) and $P_{TET}$-*TAF12*-$FLAG_3$ in ISC12 (SDC16). Lane 5-9 IP: $P_{MAL2}$-*TAF12* (ISC12), *TAF12*-$FLAG_3$ (SKC6), $P_{TET}$-*TAF12L*-$FLAG_3$ in ISC12 (SDC15), $P_{TET}$-*TAF12*-$FLAG_3$ in ISC12 (SDC16) and *TAF12L*-$FLAG_3$ (SKC3) respectively.

**Figure S3. Ectopically *$P_{TET}$* expressed Histone fold domains of CaTaf12 variants complement the absence of the full-length protein. (A-C)** Western blot confirmation of *$P_{TET}$* expressed HFD-*TAF12L* and HFD-*TAF12* complemented and cross-complemented strains**.** All the strains were grown in YPD+Doxycycline (50μg/ml) media for 7h. Whole cell extracts were made, run on 10-15% SDS PAGE gel and blotted to nitrocellulose membranes. The membranes were probed with anti-FLAG (1:1000) and anti-G6PDH (1:3000) antibodies (A) Lane 1: WT untagged (SN95), lane 2: *TAF12*-$FLAG_3$ in WT (SKC6), lane 3: $P_{MAL2}$-*TAF12* (ISC12), lane 4-5: two biological replicates for $P_{TET}$ expressed HFD*TAF12*-$FLAG_3$ in ISC12 (SDC5) and lanes 6-7: two biological replicates for $P_{TET}$ expressed HFD*TAF12L*-$FLAG_3$ in ISC12 (SDC8). (B) Lane 1: WT untagged (SN95), lane 2: *TAF12L*-$FLAG_3$ in WT (SKC3), lane 3: $P_{MAL2}$-*TAF12L* (ISC11), lane 4-5: two biological replicates for $P_{TET}$ expressed HFD*TAF12*-$FLAG_3$ in ISC11 (SDC2) and lanes 6-7: two biological replicates for $P_{TET}$ expressed HFD*TAF12L*-$FLAG_3$ in ISC11 (SDC7). (C) Lane 1: *TAF12L*-$FLAG_3$ in WT (SKC3), lane 2: WT untagged (SN95), lane 3: $P_{MAL2}$-*TAF12L* (ISC11), lane 4: *taf12lΔ/Δ* (ISC36), lane 5-6: two biological replicates for $P_{TET}$ expressed HFD*TAF12L*-$FLAG_3$ in ISC36 (SDC10) and lanes 7-8: two biological replicates for $P_{TET}$ expressed HFD*TAF12*-$FLAG_3$ in ISC36 (SDC11). Asterisk (*) represents non-specific bands. **(D)** Effect of ectopic expression of Histone fold domains (HFDs) of *TAF12* and *TAF12L* from *TET* promoter on growth of *taf12lΔ/Δ* mutant. WT strain (SN95), *taf12lΔ/Δ* (ISC36), $P_{TET}$-*HFDTAF12L*-$FLAG_3$ in *taf12lΔ/Δ* (SDC10), $P_{TET}$-*HFDTAF12*-$FLAG_3$ in *taf12lΔ/Δ* (SDC11) and vector only control (SDC12) was grown in YPD liquid media till saturation and then serial dilutions were spotted on YPD, YPD+Doxycycline (50μg/ml), YPD+menadione (90μM) and YPD+Doxycycline+menadione plates.

**Figure S4. Histone fold domain of Taf12 can interact with both Taf4 and Ada1 proteins in vitro. (A)** GST-Ada1 specifically pull down HFDTaf12L. Equal volume of Glutathione Sepharose 4B beads was incubated with 1nmole, 2nmole and 4nmole of GST-Taf4 (Lanes 6-9), GST-Ada1 (Lanes 11-14) and GST (Lanes 15-18). Beads were washed and incubated with 1nmole of purified HFDTaf12L-$His_6$. Beads were boiled and resolved on 4-15% SDS-PAGE gel. Lane 1: input for GST-Taf4, Lane 2: input for GST-Ada1, Lanes 3-5: three dilutions of input for HFDTaf12L-$His_6$, lanes 4, 8 and 15 represents control GST-Taf4, GST-Ada1 and GST bound beads respectively. **(B)** Quantification of amount of HFDTaf12L from (A) being pulled down with different amounts of GST-Ada1, GST-Taf4 and GST. The y-axis represents the percentage amount of HFDTaf12L pulled down normalized to the amount used as input. **(C)** Both GST-Taf4 and GST-Ada1 can pull down HFDTaf12. Equal volume of Glutathione Sepharose 4B beads was incubated with 1nmole, 2nmole and 4nmole of GST-Taf4 (Lanes 6-9), GST-Ada1 (Lanes 11-14) and GST (Lanes 15-18). Beads were washed and incubated with 1nmole of purified HFDTaf12-$His_6$. Beads were boiled and resolved on 4-15% SDS-PAGE gel. Lane 1: input for GST-Taf4, Lane 2: input for GST-Ada1, Lanes 3-5: three dilutions of input for HFDTaf12-$His_6$, lanes 4, 8 and 15 represents GST-HFDTaf4, GST-HFDAda1 and GST bound beads respectively. **(D)** Quantification of amount of HFDTaf12-$His_6$ from (C) being pulled down with different amounts of GST-HFDAda1, GST-HFDTaf4 and GST. The y-axis represents the percentage amount of HFDTaf12L-$His_6$ pulled down normalized to the amount used as input for pull down. **(E)** GST-Ada1 interaction with HFD-Taf12L is more selective than interaction with HFD-Taf12. Equal volume of Glutathione Sepharose 4B beads was incubated with 1nmole of GST-Taf4 (Lanes 6-9), GST-Ada1 (Lanes 11-14) and GST (Lanes 15-18). Beads were washed and incubated with 1nmole of purified HFDTaf12L-$His_6$ (lanes 8, 13 and 17), 1nmole of purified HFDTaf12-$His_6$ (lanes 9, 14 and 18) and with a mixed sample containing 1nmole each of HFDTaf12L-$His_6$ and HFDTaf12-$His_6$ (lanes 7, 12 and 16). Beads were boiled and resolved on 4-15% SDS-PAGE gel. Lane 1: input for GST-Taf4, Lane 2: input for GST-Ada1, Lanes 3-5: three dilutions of input for HFDTaf12-$His_6$, lanes 6, 11 and 15 represents control GST-Taf4, GST-Ada1 and GST bound beads respectively. **(F)** Quantification of amount of HFDTaf12L and HFDTaf12 being pulled down (from E) with GST-Ada1, GST-Taf4 and GST respectively. The y-axis represents the percentage amount of HFDTaf12L-$His_6$ and HFDTaf12L-$His_6$ pulled down normalized to the amount used as input. Statistical significance for each histogram was calculated by student T-test.

**Figure S5. CaTaf12 variants contain a highly conserved C-terminal Histone fold domain (HFD) but only CaTaf12L contains Tra1 interacting region. (A)** Schematic representation of domain architecture of CaTaf12 variants and yTaf12. **(B)** Multiple sequence alignment of only the Tra1 Interacting regions (TIR) of CaTaf12 variants and yTaf12. The different protein sequences were aligned by MUSCLE in Jalview. The conserved TIR residues (Wang et al., 2020) are highlighted as boxed residues. Asterisk (*) represents critical residue. **(C)** Multiple sequence alignment of histone fold domain of Taf12 proteins of *Saccharomycotina*. MSA of C-terminal conserved region of 35 Taf12 protein sequences of *Candida spp., Saccharomyces spp., Schizosaccharomyces pombe, Arabidopsis thaliana, Drosophila melanogaster, Mus musculus* and *Homo sapiens.* The different protein sequences were aligned by MUSCLE in Jalview. The experimentally described secondary structure is shown below the alignment. The residues shown earlier to make the hydrophobic interaction interface in hTAF12-hTAF4 crystal structure (Werten et. al., 2002) are highlighted as boxed residues. The alphabetical symbol, h represents hydrophobic residues, s represents small sized residues and A represents Alanine.

**Figure S6. CaTaf12 variants Histone fold domains (HFD) assume similar conformations. (A i-iii)** Alpha fold model of Histone fold domains (HFDs) of CaTaf12 variants. (i) HFD of Taf12L (ii) HFD of Taf12 and (iii) superimposition of alpha-fold model of HFDTaf12L and HFDTaf12. The amino-acid residues specific to HFDTaf12L or HFDTaf12 like sequences from *Candida* clade species are labeled and shown in gray color. **(B i-iii)** Alpha fold model of Histone fold domains (HFDs) of CaTaf4 and CaAda1. (i) HFD of Ada1 (ii) HFD of Taf4 and (iii) superimposition of alpha-fold model of HFDAda1 and HFDTaf4.

**Figure S7. Helix swap chimera constructs of *CaTAF12* variants show robust expression from *TET* promoter. (A-C)** Western blot confirmation of $P_{TET}$ expressed Taf12L and Taf12 chimeric constructs in $P_{MAL2}$-*TAF12* (ISC12) strain. All the strains were grown for 7h in YPD+doxycycline liquid media at 30°C and 220rpm and whole cell extracts were made, run on 10% SDS PAGE gel and blotted to nitrocellulose membranes. The membranes were probed with anti-FLAG (1:3000) and anti-G6PDH (1:3000). **(A)** Lane 1: $P_{MAL2}$-*TAF12* (ISC12), lane 2: *TAF12*-$FLAG_3$ in WT (SKC6), lane 3: $P_{TET}$ expressed *TAF12LC1*-$FLAG_3$ in ISC12 (SDC41), lane 4: $P_{TET}$ expressed *TAF12LC2*-$FLAG_3$ in ISC12 (SDC42), lanes 5-6: two biological replicates for $P_{TET}$ expressed *TAF12LC3*-$FLAG_3$ in ISC12 (SDC43), lane 7: $P_{TET}$ expressed *TAF12LC5*-$FLAG_3$ in ISC12

(SDC55), lane 8-9: two biological replicates for $P_{TET}$ expressed *TAF12C5*-$FLAG_3$ in ISC12 (SDC44). **(B)** Lane 1-2: $P_{MAL2}$-*TAF12* (ISC12), lane 3: $P_{TET}$ expressed *TAF12C4*-$FLAG_3$ in ISC12 (SDC22), lane 4: $P_{TET}$ expressed *TAF12LC4*-$FLAG_3$ in ISC12 (SDC21), lanes 5-6: two biological replicates for $P_{TET}$ expressed *TAF12C2*-$FLAG_3$ in ISC12 (SDC54). **(C)** Lane 1-2: two biological replicates for $P_{TET}$ expressed *TAF12C1*-$FLAG_3$ in ISC12 (SDC51), lane 3-4: two biological replicates for $P_{TET}$ expressed *TAF12C3*-$FLAG_3$ in ISC12 (SDC52). Asterisk (*) represents Non-specific bands.

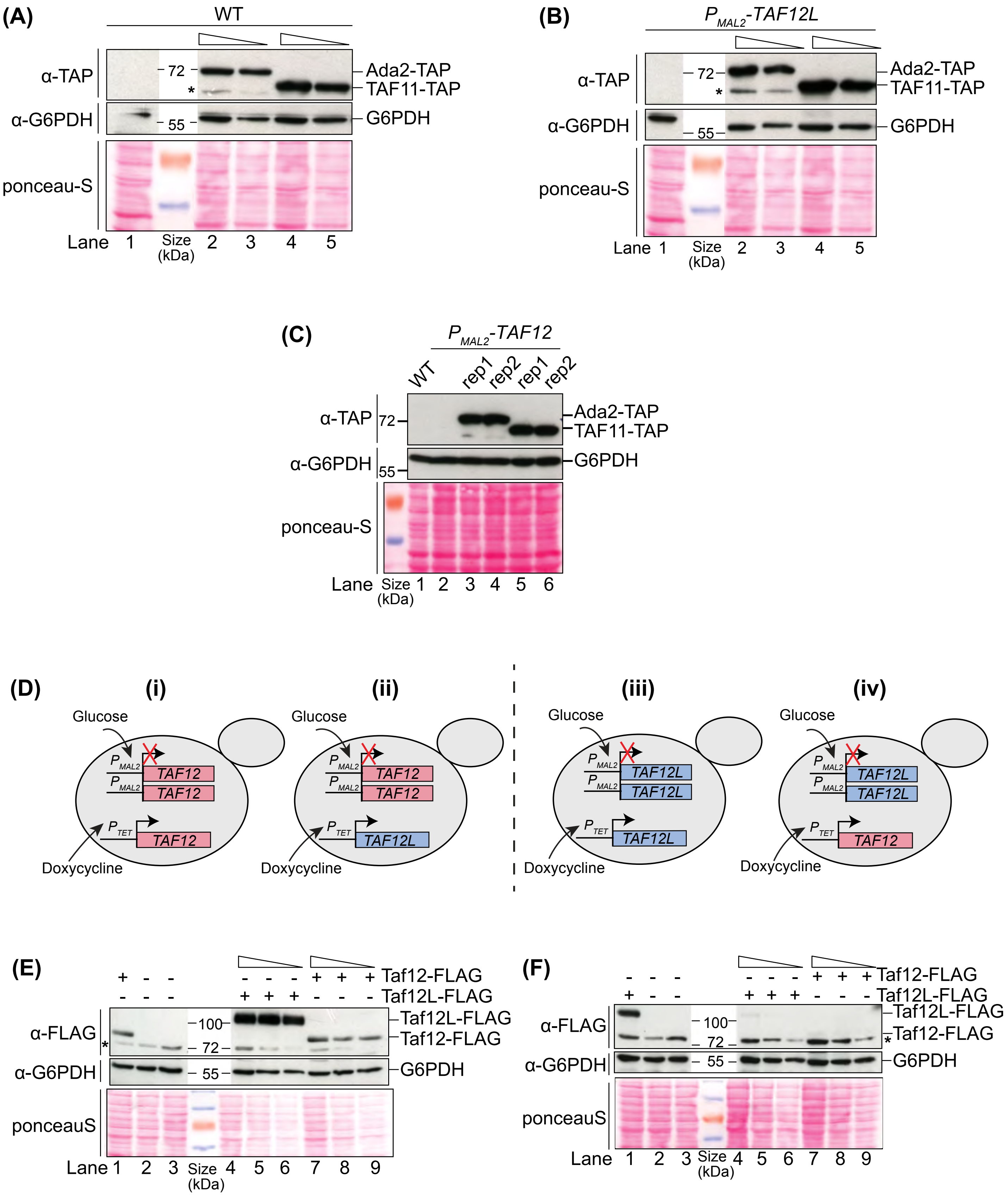
(A)
WT
α-TAP
72
*
Ada2-TAP
TAF11-TAP
α-G6PDH
55
G6PDH
ponceau-S
Lane 1 Size (kDa) 2 3 4 5
(B)
PMAL2-TAF12L
α-TAP
72
*
Ada2-TAP
TAF11-TAP
α-G6PDH
55
G6PDH
ponceau-S
Lane 1 Size (kDa) 2 3 4 5
(C)
PMAL2-TAF12
WT rep1 rep2 rep1 rep2
α-TAP
72
Ada2-TAP
TAF11-TAP
α-G6PDH
55
G6PDH
ponceau-S
Lane Size (kDa) 1 2 3 4 5 6
(D)
(i)
(ii)
(iii)
(iv)
Glucose
PMAL2
TAF12
TAF12L
PTET
Doxycycline
(E)
Taf12-FLAG
Taf12L-FLAG
α-FLAG
*
100
72
Taf12L-FLAG
Taf12-FLAG
α-G6PDH
55
G6PDH
ponceauS
Lane 1 2 3 Size (kDa) 4 5 6 7 8 9
(F)
Taf12-FLAG
Taf12L-FLAG
α-FLAG
100
72
Taf12L-FLAG
Taf12-FLAG
*
α-G6PDH
55
G6PDH
ponceauS
Lane 1 2 3 Size (kDa) 4 5 6 7 8 9

**Figure S1**

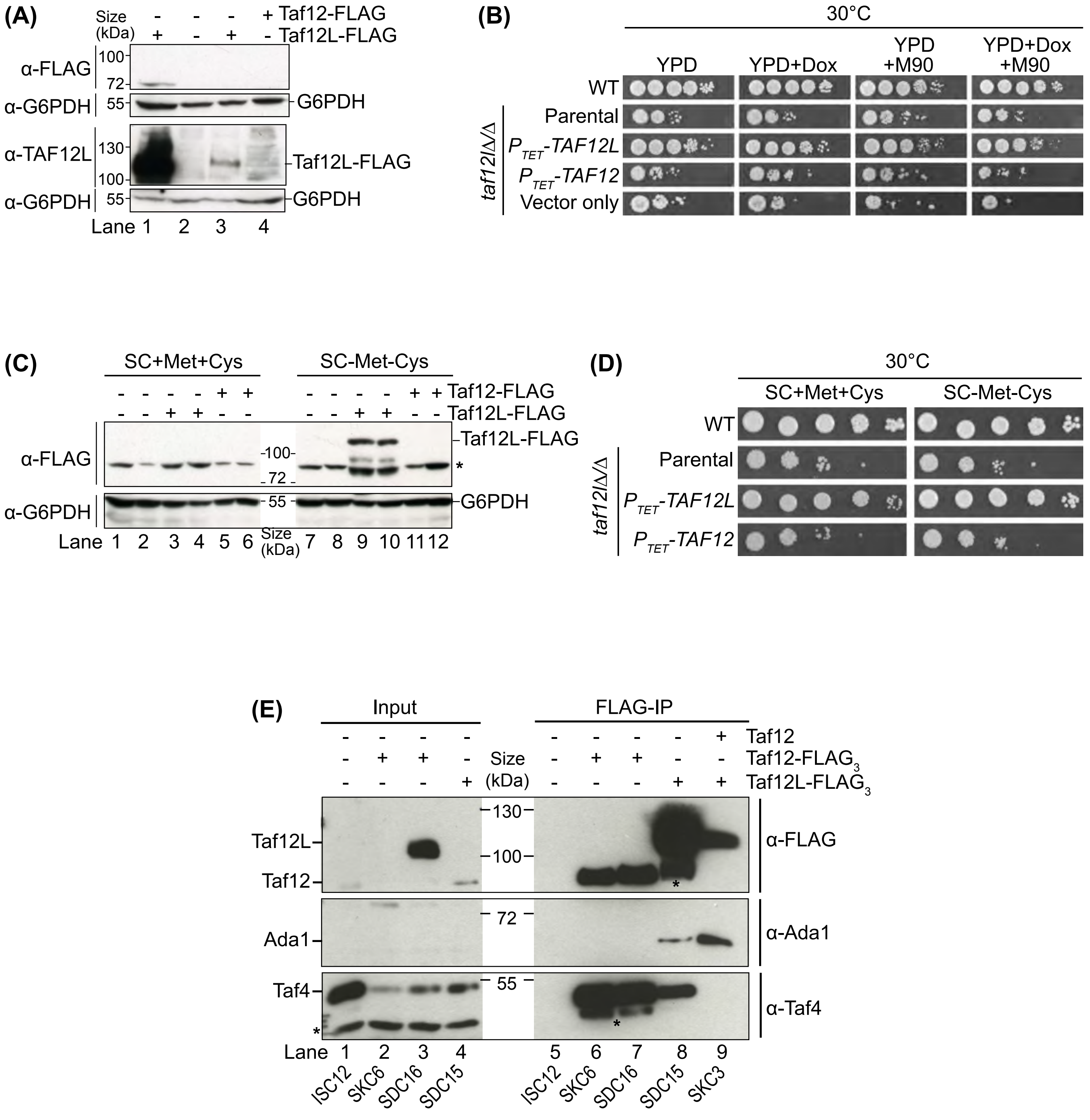
(A)
Size (kDa)
Taf12-FLAG
Taf12L-FLAG
α-FLAG
α-G6PDH
α-TAF12L
G6PDH
Taf12L-FLAG
Lane 1 2 3 4
(B)
30°C
YPD
YPD+Dox
YPD +M90
YPD+Dox +M90
WT
Parental
PTET-TAF12L
PTET-TAF12
Vector only
taf12lΔ/Δ
(C)
SC+Met+Cys
SC-Met-Cys
Taf12-FLAG
Taf12L-FLAG
α-FLAG
α-G6PDH
Taf12L-FLAG
G6PDH
Lane 1 2 3 4 5 6 Size (kDa) 7 8 9 10 11 12
(D)
30°C
SC+Met+Cys
SC-Met-Cys
WT
Parental
PTET-TAF12L
PTET-TAF12
taf12lΔ/Δ
(E)
Input
FLAG-IP
Taf12
Taf12-FLAG3
Taf12L-FLAG3
Size (kDa)
Taf12L
Taf12
Ada1
Taf4
α-FLAG
α-Ada1
α-Taf4
Lane 1 2 3 4 5 6 7 8 9
ISC12 SKC6 SDC16 SDC15
ISC12 SKC6 SDC16 SDC15 SKC3

**Figure S2**

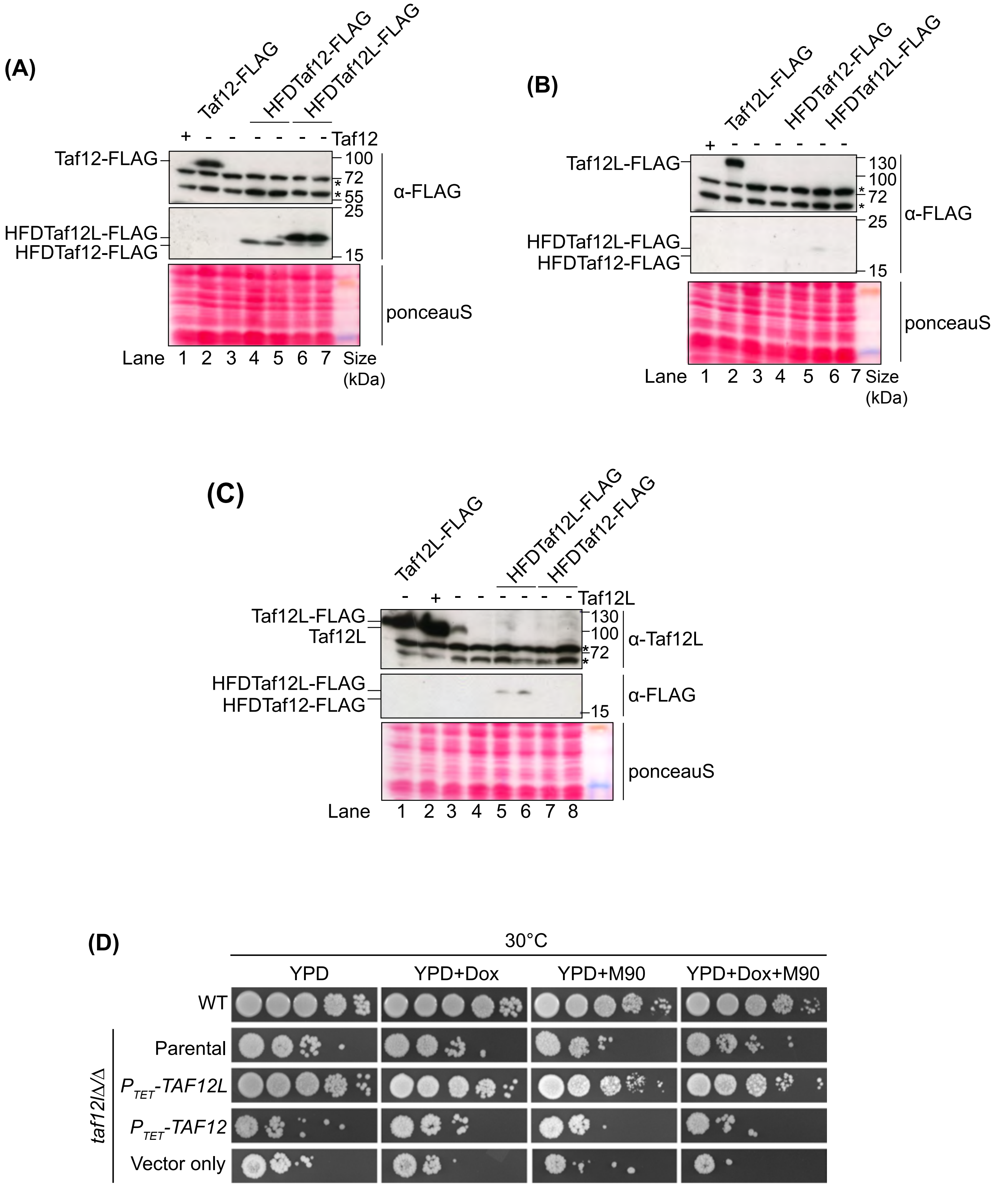
(A)
Taf12-FLAG
HFDTaf12-FLAG
HFDTaf12L-FLAG
+ - - - - - - Taf12
Taf12-FLAG
HFDTaf12L-FLAG
HFDTaf12-FLAG
100
72
55
25
15
α-FLAG
ponceauS
Lane 1 2 3 4 5 6 7 Size (kDa)
(B)
Taf12L-FLAG
HFDTaf12-FLAG
HFDTaf12L-FLAG
+ - - - - - -
Taf12L-FLAG
HFDTaf12L-FLAG
HFDTaf12-FLAG
130
100
72
25
15
α-FLAG
ponceauS
Lane 1 2 3 4 5 6 7 Size (kDa)
(C)
Taf12L-FLAG
HFDTaf12L-FLAG
HFDTaf12-FLAG
- + - - - - - - Taf12L
Taf12L-FLAG
Taf12L
130
100
72
α-Taf12L
HFDTaf12L-FLAG
HFDTaf12-FLAG
15
α-FLAG
ponceauS
Lane 1 2 3 4 5 6 7 8
(D)
30°C
YPD
YPD+Dox
YPD+M90
YPD+Dox+M90
WT
taf12l∆/∆
Parental
P_TET-TAF12L
P_TET-TAF12
Vector only

### Figure S3

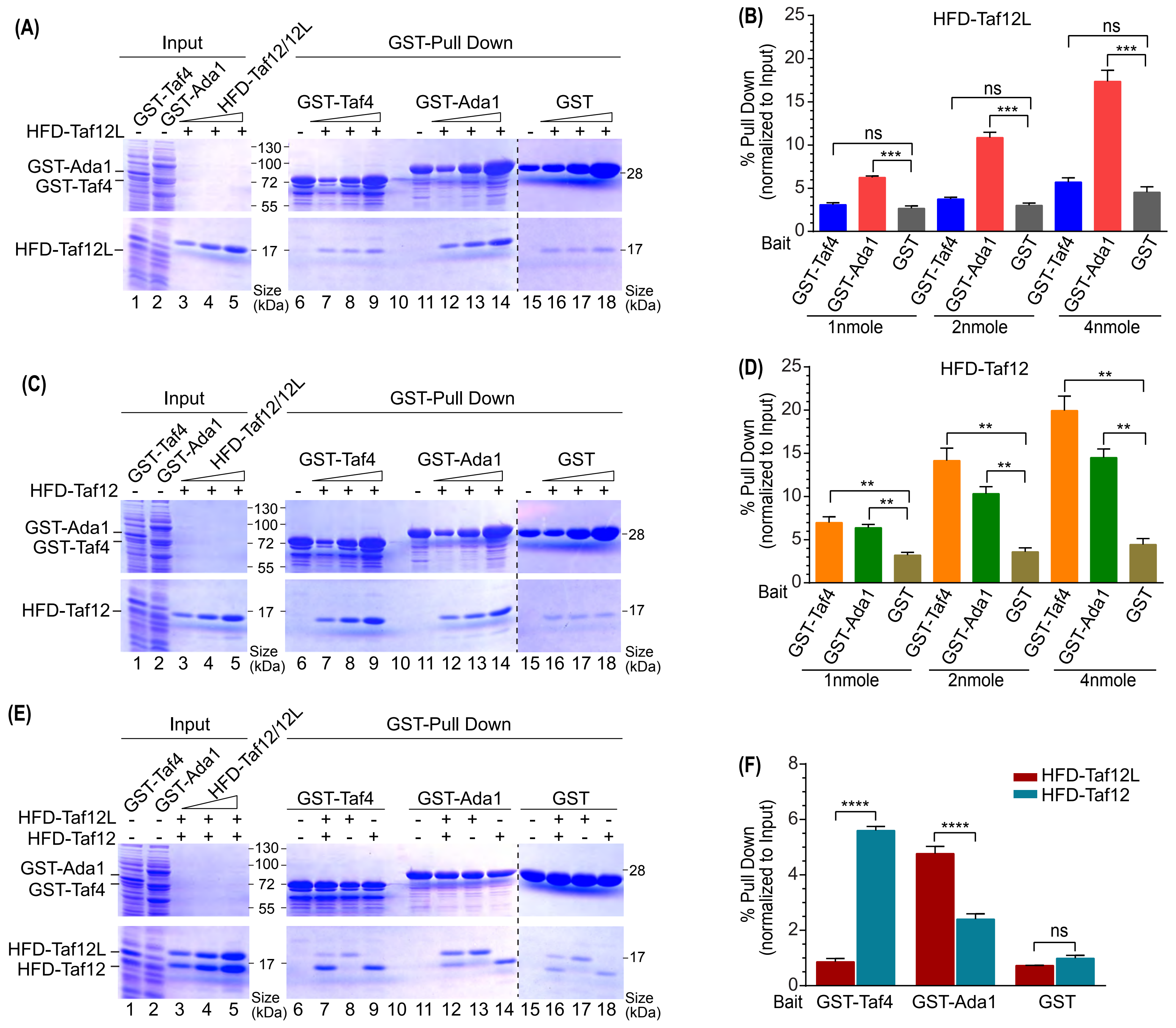
(A)
Input
GST-Pull Down
GST-Taf4
GST-Ada1
HFD-Taf12/12L
HFD-Taf12L
GST-Ada1
GST-Taf4
HFD-Taf12L
130
100
72
55
17
28
Size (kDa)
1 2 3 4 5 6 7 8 9 10 11 12 13 14 15 16 17 18
(B)
HFD-Taf12L
% Pull Down (normalized to Input)
Bait
ns
***
1nmole
2nmole
4nmole
(C)
HFD-Taf12
(D)
**
(E)
(F)
HFD-Taf12L
HFD-Taf12
****
GST


Figure S4

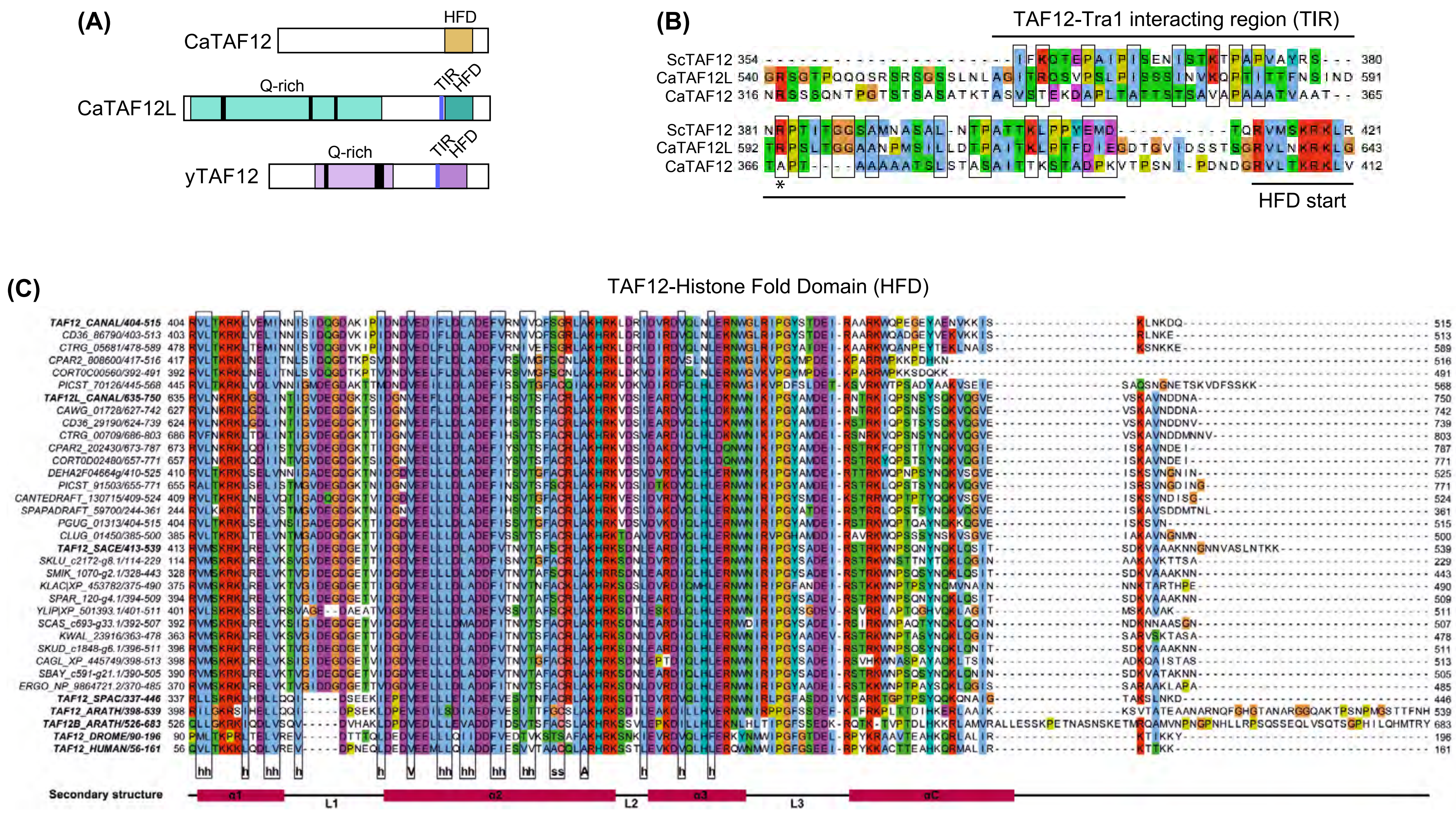
(A)
HFD
CaTAF12
Q-rich
TIR
HFD
CaTAF12L
Q-rich
TIR
HFD
yTAF12
(B)
TAF12-Tra1 interacting region (TIR)
ScTAF12 354
CaTAF12L 540
CaTAF12 316
380
591
365
ScTAF12 381
CaTAF12L 592
CaTAF12 366
421
643
412
*
HFD start
(C)
TAF12-Histone Fold Domain (HFD)
TAF12_CANAL/404-515
TAF12L_CANAL/635-750
TAF12_SACE/413-539
TAF12_SPAC/337-446
TAF12_ARATH/398-539
TAF12B_ARATH/526-683
TAF12_DROME/90-196
TAF12_HUMAN/56-161
Secondary structure
α1
L1
α2
L2
α3
L3
αC

Figure S5

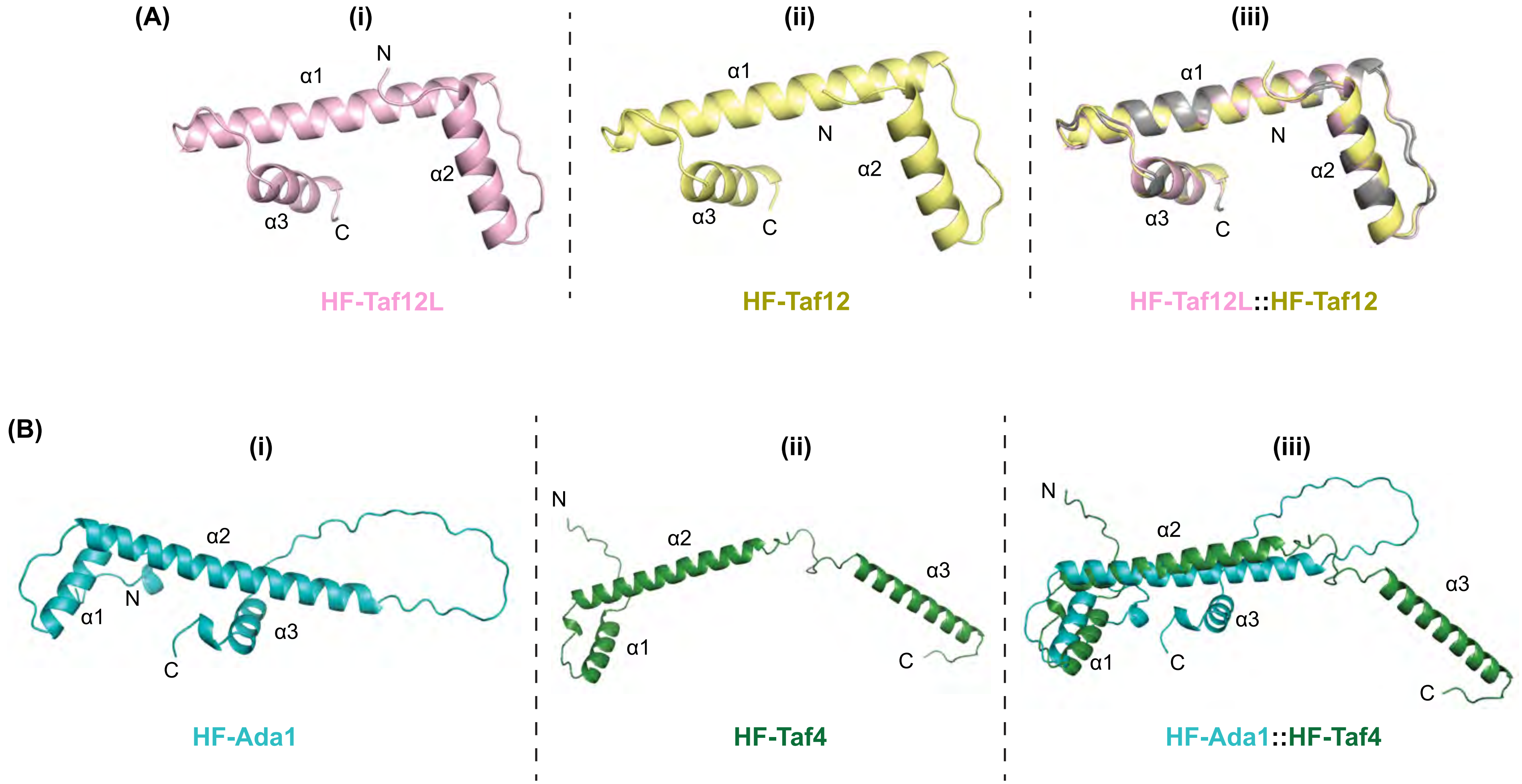
(A)
(i)
N
α1
α2
α3
C
HF-Taf12L
(ii)
α1
N
α2
α3
C
HF-Taf12
(iii)
α1
N
α2
α3
C
HF-Taf12L::HF-Taf12
(B)
(i)
α2
N
α1
α3
C
HF-Ada1
(ii)
N
α2
α3
α1
C
HF-Taf4
(iii)
N
α2
α3
α1
C
C
HF-Ada1::HF-Taf4

**Figure S6**

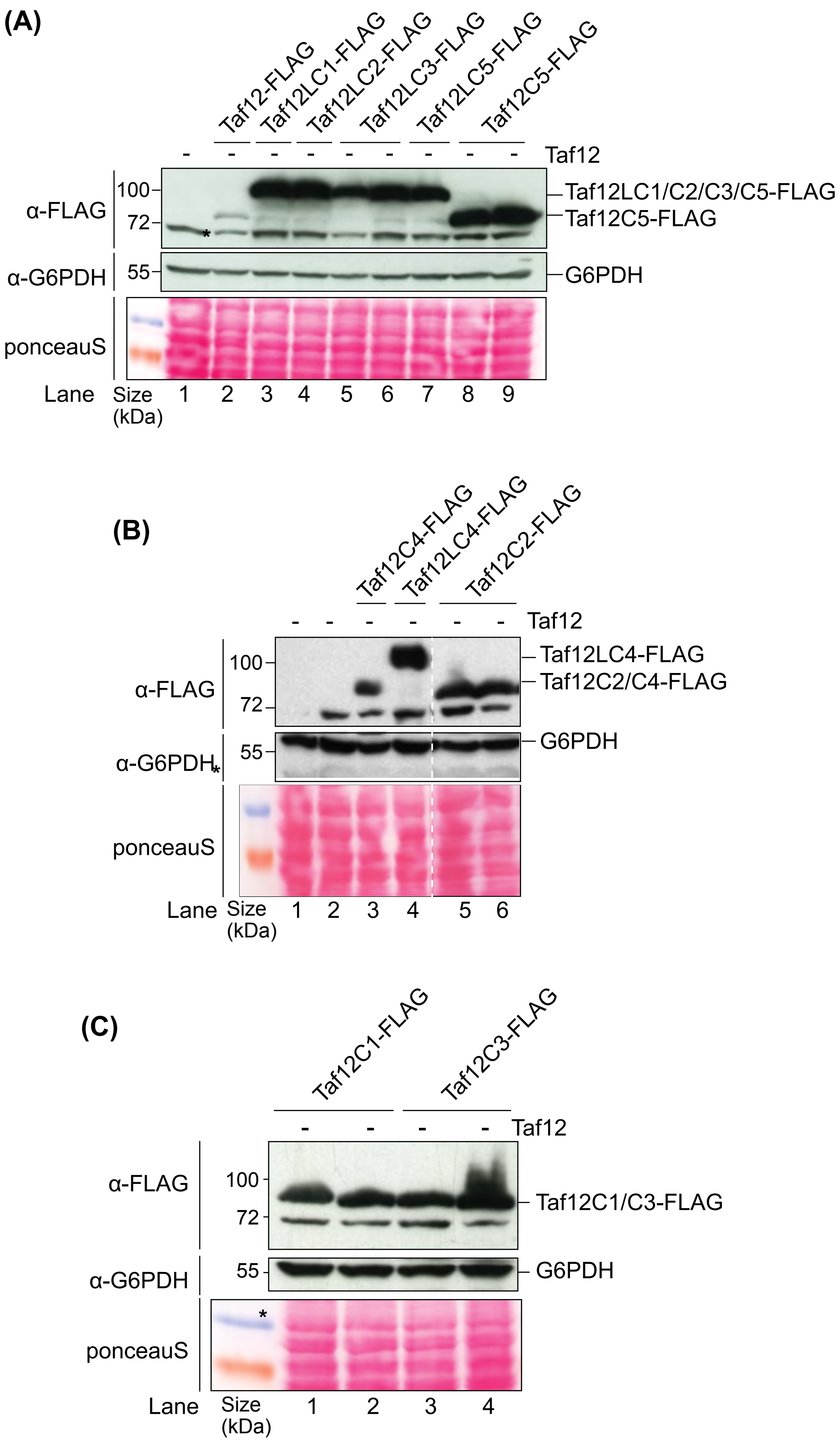
(A)
Taf12-FLAG
Taf12LC1-FLAG
Taf12LC2-FLAG
Taf12LC3-FLAG
Taf12LC5-FLAG
Taf12C5-FLAG
Taf12
α-FLAG
100
72
Taf12LC1/C2/C3/C5-FLAG
Taf12C5-FLAG
α-G6PDH
55
G6PDH
ponceauS
Lane
Size (kDa)
1 2 3 4 5 6 7 8 9
(B)
Taf12C4-FLAG
Taf12LC4-FLAG
Taf12C2-FLAG
Taf12
α-FLAG
100
72
Taf12LC4-FLAG
Taf12C2/C4-FLAG
α-G6PDH
55
G6PDH
ponceauS
Lane
Size (kDa)
1 2 3 4 5 6
(C)
Taf12C1-FLAG
Taf12C3-FLAG
Taf12
α-FLAG
100
72
Taf12C1/C3-FLAG
α-G6PDH
55
G6PDH
ponceauS
Lane
Size (kDa)
1 2 3 4

**Figure S7**
